# Plucking a string or playing a G? Choice type impacts human reinforcement learning

**DOI:** 10.1101/2021.08.25.457707

**Authors:** Milena Rmus, Amy Zou, Anne G.E. Collins

**Affiliations:** Department of Psychology, University of California, Berkeley; Helen Wills Neuroscience Institute, University of California, Berkeley, California 94720, USA

## Abstract

In reinforcement learning (RL) experiments, participants learn to make rewarding choices in response to different stimuli; RL models use outcomes to estimate stimulus-response values which change incrementally. RL models consider any response type indiscriminately, ranging from more concretely defined motor choices (e.g. pressing a key with the index finger), to more general choices that can be executed in a number of ways (e.g. getting dinner at the restaurant). But does the learning process vary as a function of the choice type? In Experiment 1, we show that it does: participants were slower and less accurate in learning correct choices of a general format compared to learning more concrete, motor actions. Using computational modeling, we show that two mechanisms contribute to this. First, there was evidence of irrelevant credit assignment: the values of motor actions interfered with the values of other choice dimensions, resulting in more incorrect choices when the correct response is not defined by a single motor action; second, information integration for relevant general choices was slower. In Experiment 2, we replicated and further extended the findings from Experiment 1, by showing that slowed learning was attributable to weaker working memory use, rather than slowed RL learning. In both experiments we ruled out the explanation that the difference in performance between two condition types was driven by difficulty/different levels of complexity. We conclude that defining a more abstract choice space used by multiple learning systems for credit assignment recruits executive resources, limiting how much such processes then contribute to fast learning.

## 1 Introduction

The ability to learn rewarding choices from non-rewarding ones lies at the core of successful goal-directed behavior. But what counts as a choice? When a child tries the pink yogurt in the left cup, and the white yogurt in the right cup, then prefers the right cup, what choice should they credit this rewarding outcome to? In their next decision, should they repeat their previously rewarding reach to the yogurt on the right, independently of its color, or should they figure out where the white yogurt is before reaching for it? Selecting the type of yogurt is a more abstract choice: it requires subsequently paying attention to the other dimension (where is the white yogurt?) and applying the appropriate motor program to execute the choice. Thus, making the more abstract choice additionally involves less abstract choices, but in this case, only the abstract choice should be credited for the yogurt’s tastiness. Knowing the relevant dimension of choice to assign credit to is essential when learning. How does choice type impact how we learn?

The theoretical framework of reinforcement learning (RL) is highly successful for studying reward-based learning and credit assignment (Sutton & Barto, 2018). However, RL as a computational model of cognition typically assumes a given action space defined by the modeler, which provides the relevant dimensions of the choice space (i.e. either the yogurt color or the cup position) - there is no ambiguity in what choices are (i.e. color such as pink/white, or side such as left/right), and the nature of the choice space does not matter (Rmus, McDougle, & Collins, 2020). As such, RL experiments in psychology tend to not consider the type of choices (a single motor-action such as pressing a key with the index finger; (A. G. E. Collins, Ciullo, Frank, & Badre, 2017; Tai, Lee, Benavidez, Bonci, & Wilbrecht, 2012), or the more general selection of a goal stimulus that is not tied to a specific motor action (Foerde & Shohamy, 2011; Daw, Gershman, Seymour, Dayan, & Dolan, 2011; Frank, Moustafa, Haughey, Curran, & Hutchison, 2007)) as important, and researchers use the same models and generalize findings across choice types. Recent research has shed some light into how participants might identify relevant dimensions of the state and choice space (Niv, 2019; Farashahi, Rowe, Aslami, Lee, & Soltani, 2017); however, this research does not address how learning occurs when the learner knows the relevant choice space but multiple dimensions of choice are nonetheless available, such as in our yogurt example.

Examining learning of responses when multiple choice dimensions may be relevant is important, however, as most of our choices in everyday life are ambiguous: did I pick the white yogurt or the one of the left? In some cases, these dimensions are hierarchically interdependent: choices can be represented at multiple levels of abstraction (e.g. have breakfast; have yogurt; have pink yogurt; have the yogurt on the right; reach for the yogurt on the right side, etc.). In such cases, a choice along a relevant dimension (yogurt color) requires a subsequent choice on a reward-irrelevant dimension (position/motor action), which thus needs to be considered for the choice’s execution, but not credited during learning. By contrast, in other cases, some choice dimensions may neither be relevant for learning nor for executing the choice – for example, the child should learn to fully ignore the color of the plate the yogurt is on for both their choice and their credit assignment.

Different types of choices may recruit different cognitive/neural mechanisms (Rescorla & Solomon, 1967). For example, previous animal models of decision-making suggest that the orbitofrontal cortex and the anterior cingulate cortex index choice outcomes for goal stimulus choices and motor action choices respectively (Luk & Wallis, 2013). Ventral striatum lesions in monkeys impaired learning to choose between rewarding stimuli, but not between rewarding motor actions (Rothenhoefer et al., 2017). In humans, recent behavioral evidence suggests that the credit assignment process is what differentiates learning more relevant choice dimensions from less relevant (here motor) ones (McDougle et al., 2016), and that there might be a hierarchical gradation of choices in terms of credit assignment. In particular, while people are capable of learning the value of both abstract rule choices and concrete action choices in parallel (Eckstein, Starr, & Bunge, 2019; Eckstein & Collins, 2019; Ballard, Miller, Piantadosi, Goodman, & McClure, 2018), they also seem to assign credit to more concrete actions by default when making abstract choices that need to be realized through motor actions (Shahar et al., 2019).

The brain relies on multiple neuro-cognitive systems for decision-making; whether choice format impacts learning similarly across systems remains unexplored. Specifically, while RL models provide a useful formalism of learning, they do not easily relate to underlying processes. Indeed, RL models are known to summarize multiple processes that contribute to learning jointly (Eckstein, Wilbrecht, & Collins, 2021), such as the brain’s RL mechanism, but also episodic memory (Wimmer & Shohamy, 2012; Bornstein & Daw, 2013; Vikbladh et al., 2019; Bornstein, Khaw, Shohamy, & Daw, 2017; Poldrack et al., 2001), or executive functions (Rmus et al., 2020; A. G. E. Collins & Frank, 2012). Here we focus on working memory (WM), which has also been shown to contribute to learning alongside RL (A. G. E. Collins & Frank, 2012, 2018; A. G. E. Collins et al., 2017). If choice type matters for learning, does it matter equally for each cognitive system that contributes to learning, or differently so?

In summary, there is a two-fold gap in our understanding of how choice format impacts learning. First, when multiple choice dimensions are available, does the relevant choice type impact learning, and if so, through what computational mechanisms? We consider, in particular, the important case where one relevant choice dimension needs to be executed through a second, irrelevant choice dimension (a motor action); and how this contrasts to learning when one dimension is fully irrelevant to both choice and learning. Second, are the differences rooted in the brain’s RL system, WM, or both? To address this gap, we designed a task that directly compares learning to make choices along two orthogonal dimensions, with different levels of generality or interdependence, when there is no ambiguity about which choices are relevant to the learning problem. In our task, one choice dimension is a spatial position that directly maps on to a consistent motor actions, and the other is a more general choice dimension, conceptualized as the selection of stimulus goals that constrain a downstream selection of an overall irrelevant spatial position and corresponding motor action. In a second experiment, we manipulated learning load to separately identify WM and RL contributions to learning, and investigated with computational modeling how choice matters in both systems.

Our results across two experiments suggest that choice type impacted learning strongly, resulting in slower learning when the relevant choice dimension was more general and required execution along another dimension. This was in part driven by an incorrect, asymmetric credit assignment to less general choices when they were irrelevant. Further-more, WM (rather than RL) mechanisms seemed to drive the deficits in performance in the more general choice format condition, indicating that defining a more general action space, shared by multiple choice systems, recruited limited executive resources. In both experiments, we ruled out the simple explanation that the performance difference was driven by an effect of difficulty by 1) implementing experimental controls that minimize this concern, and 2) ruling out predictions of a pure difficulty effect in analyses and modeling.

## 2 Methods

### 2.1 Participants

#### 2.1.1 Experiment 1

Our sample for experiment 1 consisted of 82 participants (40 female, age mean (SD) = 20.5(1.93), age range = 18-30), recruited from the University of California, Berkeley Psychology Department’s Research Participation Program (RPP). In accordance with the UC Berkeley Institutional Review Board policy, participants provided written informed consent before taking part in the study. They received course credit for their participation. To ensure that the participants included in analyses were engaged with the task, we set up an exclusion criterion of 0.60 or greater average accuracy across all task conditions. This cutoff was determined based on an elbow point in the group’s overall accuracy in the task (Fig. 12). We excluded 20 participants based on this criterion, resulting in a total sample of 62 participants for the reported analyses.

#### 2.1.2 Experiment 2

For the second experiment, we recruited 75 participants (54 female, 1 preferred not to answer; age mean (SD) = 20.34(2.4), Age-range=18-34) from the University of California, Berkeley RPP. Participants completed the experiment online (de Leeuw, 2015), and received course credit for their participation. Using the same exclusion criteria as previous experiment (based on the distribution of average accuracy), we excluded 18 participants, resulting in the total sample of 57 participants.

### 2.2 Experimental protocol

#### 2.2.1 Experiment 1

Learning Blocks. Participants were instructed that they would be playing a card sorting game, and that on each trial they would sort a card into one of three boxes. Their goal was to use reward feedback to learn which box to sort each card in. The boxes were labeled with 3 different colors (green, blue and red), and participants chose one of the boxes by pressing one of three contiguous keyboard keys (corresponding to the box position) with their index, middle and ring finger. Importantly, the color of the boxes changed positions on different trials (i.e. the blue box could appear on the right side on trial n, and in the middle on trial n+1). Participants received deterministic feedback after each selection (+1 if they selected the correct box for the current card, 0 otherwise).

Before the experiment, participants read detailed instructions and practiced each task condition. The task then consisted of 8 blocks, divided into three conditions. Each of the three conditions was defined by its distinct sorting rule. In the label condition, the correct box for a given card was defined deterministically by the box’s color label (Fig. 1A). For instance, if the blue box was the correct choice for a given card, participants were always supposed to select the blue box in response to that card, regardless of which key mapped onto the blue box on a given trial. In the position condition, the correct box was defined deterministically by the box’s position (left/middle/right). For example, the correct response of a given card would always be achieved by pressing the leftmost key with the index finger, regardless of the box color occupying the left position (Fig. 1B). The sorting rule in the position control condition was identical to the sorting rule in the position condition, but the boxes were not tagged with color labels. This condition allowed us to assess participants’ baseline performance when only one response type (e.g. position, but not the label) was available. Importantly, participants were explicitly told the sorting rule (position or label) at the beginning of each block, in order avoid any performance variability that may arise as a function of rule inference and uncertainty. Following the 8 learning blocks, participants performed two additional tasks; these are not the focus of the current paper and are not analyzed here.

**Figure 1:**
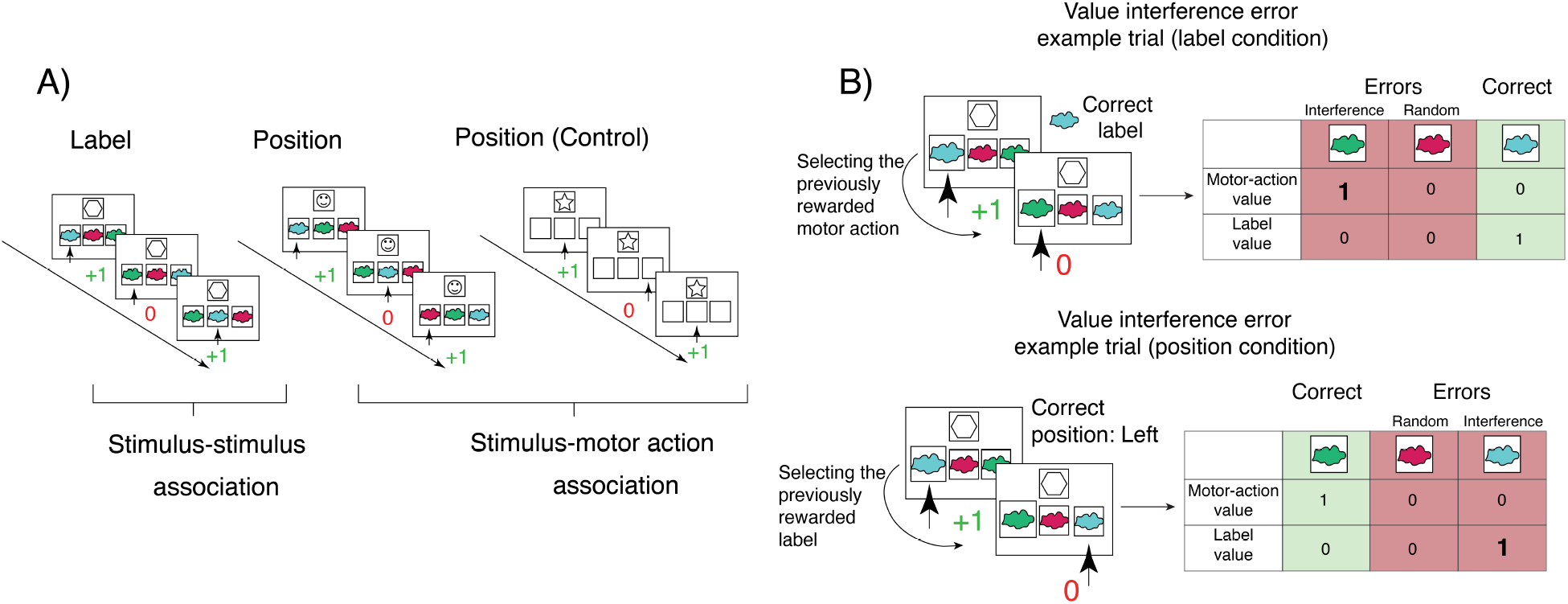
Experiment design. A) Participants played a card-sorting game with 3 different conditions: Label (learning which box color is correct for each card - more general choice), Position (learning which motor action/position is correct for each card - less general choice), Control (identical sorting rules as position condition, but without label-tagged boxes). B) We assumed that participants track card-dependent reward history for both positions and labels, and that both of these contribute to choice selection process, sometimes resulting in interference errors. Note that the card-dependent reward history is cumulative (tracked across all past trials during which the given card was presented, rather than only one-trial back), but for simplicity of illustration we only show 1-back trial in the panel B.

Out of 8 blocks in total, 2 were control condition blocks, 3 were position condition, and 3 were label condition. Block order was pseudo-randomized: participants completed a control block first and last, while blocks 2-7 conditions were randomly chosen within subjects, but counterbalanced across subjects. In each block, participants learned how to sort 6 different cards; we used a different set of images to represent cards in each block. The boxes were labeled with the same 3 colors across all blocks, except the position control blocks, where the boxes were not labeled. Participants experienced 15 repetitions of each card, resulting in 90 trials per block; trial order was pseudo-randomized to ensure a uniform distribution of delays between repetitions of the same card in a block. We controlled for the card-dependent position-label combinations across trials. Specifically, each label occurred in each position an equal number of times (i.e. the blue label occurred 5 times on the left, right and middle box for each card). We also ensured that the position-label combinations were evenly distributed across the task (i.e. blue-middle combination did not occur only during the first quarter of block trials).

Single trial structure: controlling for the difficulty difference. On each trial, participants first saw the three boxes with their color labels underneath a fixation cross at the center of the screen. After 1 second, the card appeared in the center of the screen, replacing the fixation cross. Participants were allowed to press a key only once the card appeared, with a 1-second deadline. Following their response, participants received feedback (+1 or 0) that remained on the screen for 1 second, followed by a 1 second inter-trial interval (fixation cross). This trial structure allowed participants to identify where each color label was positioned, thus minimizing a potential advantage of the position conditions, where participants did not need to know where colors were on a trial-by-trial basis in order to make a correct response. In other words, it is possible that any condition-based difference in performance might stem from the label condition being more difficult. However, giving participants time to identify where each color is positioned prior to the onset of the card decreases the difference between the conditions in terms of difficulty, making this confound less likely.

We designed the label and position condition to engage choice processes with different degrees of generality. The position condition should capture the less general choice process in which the rewarding response is defined by a single motor action, and the label is irrelevant to both choice and learning. The label condition, on the other hand, captures a more general choice process in which the rewarding response (i.e. choice of the correct label) can be made by identifying one of three positions and executing any of the three motor actions, depending on where the correct box label is positioned on the given trial - such that the other dimension (position) remains irrelevant for learning, but is relevant for choice.

#### 2.2.2 Experiment 2

The task design for Experiment 2 was the same as the task design for Experiment 1, with one important exception - we varied the number of cards per block between 2 and 5, for both position and label conditions. This manipulation has previously been shown to enable computational modeling to disentangle working memory and reinforcement learning processes (A. G. E. Collins & Frank, 2012). The order of blocks was counterbalanced across participants; participants completed either label or position blocks first, with the order of set sizes randomized for the first completed condition, and then repeated for the second. In addition, we removed the control condition, given that we previously observed no difference between position and control. Participants completed 4 blocks of position and label each, where each condition block had a different set size.

### 2.3 Analyses

### 2.4 Model-independent analyses

In addition to general diagnostics and standard statistical analyses (see results), we sought to analyze participants’ choices and RTs as a function of how often each motor action and each label had been rewarded for each card. Specifically, we computed card-dependent cumulative reward history (CRH) for both positions *P* and labels *L* on each trial for a card *C*, in each condition:

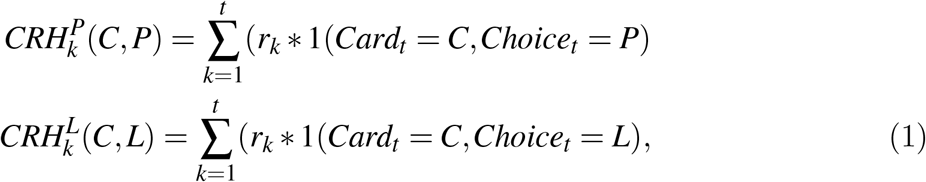

where *r*_*k*_ is the outcome at trial *k* in the block, and 1 is the indicator function taking value 1 if the card and position/label match C and P/L, and 0 otherwise. We used this metric to analyze how the integration of two value sources shaped choices when choice format was less/more general. In particular, in the example of the position condition, the position CRH for a card and its associated correct position indicated the past number of correct choices, while the CRH for other positions was 0. By contrast, in the same position condition, the label CRH for a card reflected how often each label had been rewarded due to this label being in the correct position. All label CRH values in the position condition were expected to be close to each other because label positions were counterbalanced, but slight differences due to past choice randomness could be predictive of biases in future choice. The opposite was true in the label condition.

To analyze how the value integration for each type of choice shaped decisions, we focused on the error trials and computed the proportion of errors driven by the other irrelevant choice dimension. We reasoned that if participants were lapsing randomly, any of the two possible errors should be equally likely. However, if participants experienced value interference, they should be more likely to select the error with the higher CRH in the irrelevant dimension. In the label condition, such an interference error would look like selecting the position/motor action that was rewarded on the previous trial, even though the correct label had switched positions since (Fig. 1B). In the position condition, an interference error would occur when participants selected the previously rewarded label that had switched positions, instead of the label currently corresponding to the position/motor action that is always correct for the given card (Fig. 1B).

We ran a trial-by-trial analysis using a mixed effects general linear model to characterize choices. We used trial-by-trial reward history difference *RHD* = *CRH*(*chosen*) *− mean*(*CRH*(*unchosen*)) between chosen and unchosen boxes, for both positions and labels, and tested whether this discrepancy modulated accuracy and RTs. If participants implemented an optimal decision strategy, their accuracy and RTs should increase and decrease respectively with an increased RHD in the relevant choice dimension (i.e. label RHD in label condition, position RHD in position conditon). Alternatively, contribution of the irrelevant dimension RHD (i.e. position RHD in label condition or vice versa) would serve as evidence of value interference. Our mixed effects models had the following general structure:

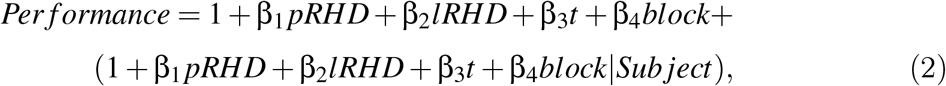

where *pRHD* is RHD based on position reward history, and *lRHD* is RHD based on label reward history. Performance can refer to either accuracy (coded as correct/incorrect) or response times.

In the analysis of Experiment 2 data, we also ran mixed-effects models including predictors indexing WM mechanisms (set size and delay between current stimulus and the most recent rewarded stimulus presentation; indexing capacity and susceptibility to decay properties of WM respectively), and RL effects (dimension-relevant, card-dependent reward history, calculated from the cumulative number of earned points for each card, indexing reward-based learning).

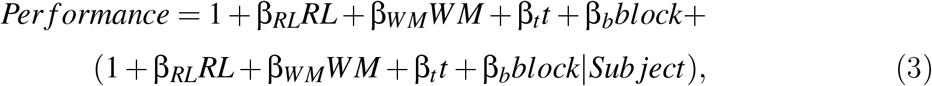

where RL corresponds to RL factors such as reward history, and WM corresponds to WM factors such as decay and set size. Note that this is a general structure to demonstrate how we structured the mixed effects model, but set size and decay were entered as separate predictors.

In other words, we explored the effects of interest on a group level, as well as how the estimates of these effects vary across individual participants. We included a predictor for trial number in this model, to ensure that reduction in RTs is not simply conflated with practice effect/task progression. In addition, we added block number as one of the regressors, in order to capture overall improvement in performance across the task.

### 2.5 Computational modeling

Reinforcement Learning-Working Memory (RL-WM): In order to computationally quantify the differences in learning processes in motor choice/general choice conditions, we used a set of hybrid reinforcement learning (RL) and working memory (WM) models. Our baseline assumption was that in the RL process, participants track and update two independent sets of stimulus-action value tables, corresponding to the two possible choice spaces: a card-position value table, and a card-label value table. We also assumed that the choice policy may reflect a mixture of both the relevant and the irrelevant value tables, potentially leading to interference errors when the value of irrelevant choice dimension (position/label) contributes to the choice process (Fig. 2A). In addition to the RL module, a WM module allows us to capture the contribution of WM to performance, particularly in the second experiment where the set size is varied between 2 and 5, since WM contribution is modulated by the cognitive load (A. G. E. Collins & Frank, 2012; A. G. Collins, 2017; A. G. E. Collins & Frank, 2018). WM also potentially tracks associations between cards and two choice types, and similarly to RL, its policy may reflect a mixture of both relevant and irrelevant associations. We investigated a range of models to pinpoint the computational mechanisms of divergence between the learning processes in the two conditions, by varying the extent to which the models allowed for condition-dependent specificity/model-parameters.

**Figure 2:**
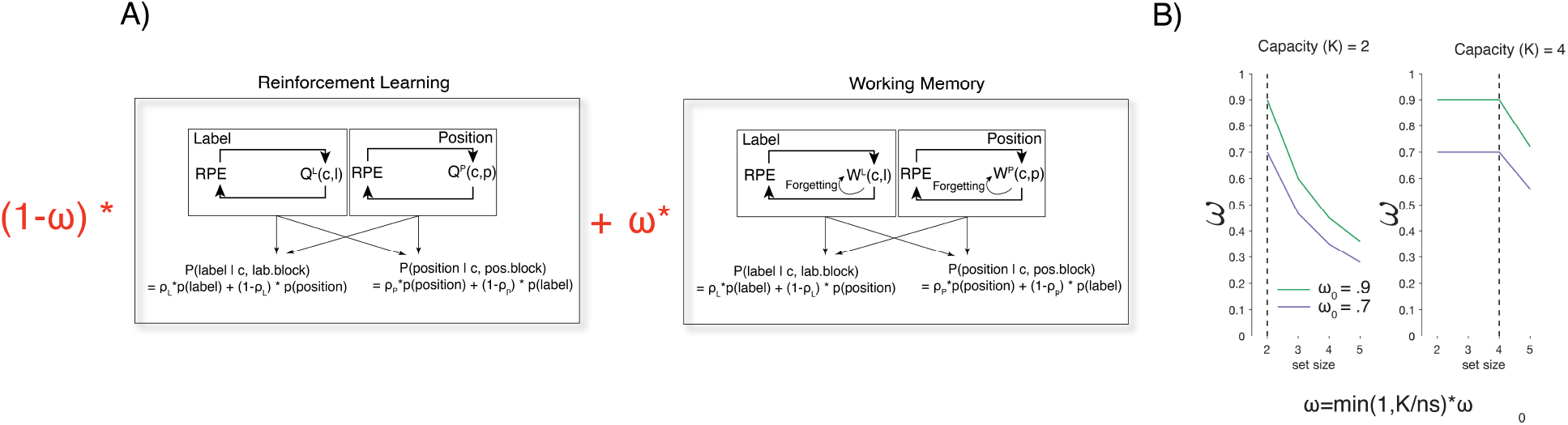
A) In experiment 1 we used RL model variants, which assume incremental, feedback-driven learning. In Experiment 2, we combined RL and WM modules, under the assumption that learning is a weighted interaction between RL and WM systems. B) The extent to which participants relied on WM was determined by the WM weight parameter(**ω**), proportional to participants’ WM capacity (K), and inversely proportional to set size.

#### 2.5.1 RL learning rule

The RL module assumes incremental learning through a simple delta rule (Sutton & Barto, 2018). Specifically, on each trial *t*, the values of labels *Q*_*L*_(*c, l*) and positions *Q*_*P*_(*c, p*) for the trial’s card *c* and chosen labels and positions *l* and *p* are updated in proportion to the reward prediction error:

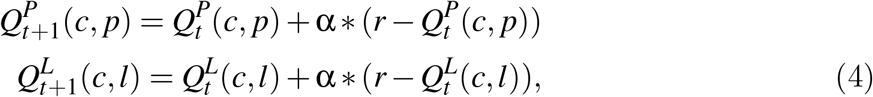

where **α** is the learning rate, and *r* = 0*/*1 is the outcome for incorrect and correct trials. *Q*-tables are initialized at 1*/*3 (3 = total number of positions/labels) at the start of each block to reflect initial reward expectation in the absence of information about new cards.

#### 2.5.2 WM learning rule

Unlike RL, WM is a one-shot learning system able to retain perfectly the previous trial’s information. This is modeled by storing the immediate outcome as the stimulus-response weight.

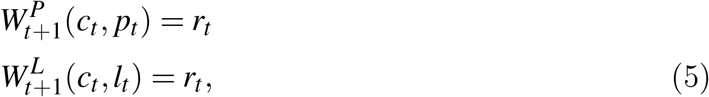

Prior work in similar tasks (Sugawara & Katahira, 2021; Frank et al., 2007; Gershman, 2015; Niv, Edlund, Dayan, & O’Doherty, 2012) has shown an asymmetry in learning based on positive/negative feedback, such that individuals are less likely to integrate negative feedback while learning rewarding responses. Thus, we included a learning bias parameter (0 *≤ LB ≤* 1), which scales the learning rate **α** by *LB* when participants observe the negative feedback. We applied *LB* to both RL and WM (for both position and label dimensions, showing only an example for position here):

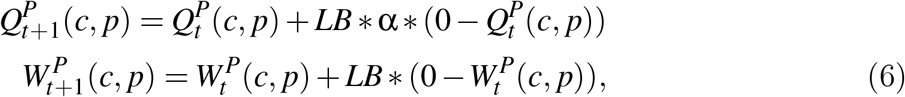

The weights stored in WM are susceptible to decay (φ) at each trial, which pulls the weights to their initial values 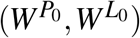:

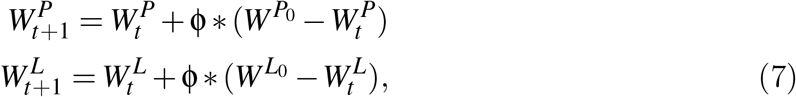

#### 2.5.3 Policy

We used the softmax function to transform WM weights and RL Q-values into choice probabilities to produce position choice policies 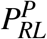 and 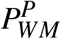:

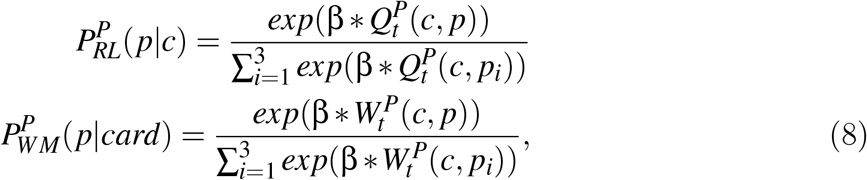

We apply the same softmax transformation to the label Q- and W-tables to obtain label choice policies 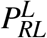 and 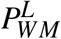. This policy permits the selection of choices with higher Q-values/weights with higher probability. The softmax β is the inverse temperature parameter, which controls how deterministic the choice process is. For each module, the overall choice policy is a mixture of both policies, determined by mixture parameters, ρ:

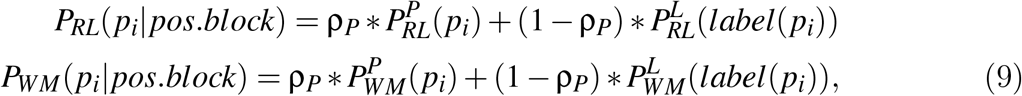

We apply the same mixture process with mixture weight ρ_*L*_ for the label dimension blocks:

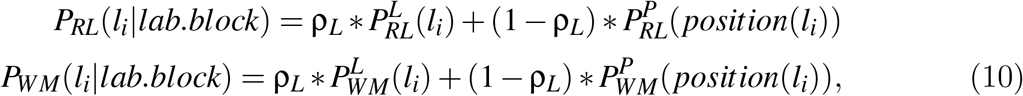

The RL-WM model posits that choice comes from a weighted mixture of RL and WM, where one’s reliance on WM is determined by the WM weight (**ω**) parameter:

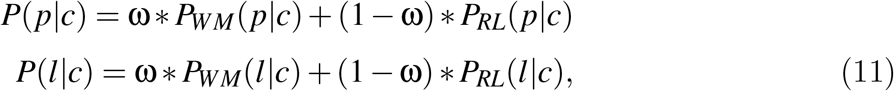

where **ω** reflects the likelihood of an item being stored in working memory and is proportional to the ratio of capacity parameter (K) and block set size, scaled by the baseline propensity to rely on WM (**ω**_0_;Fig 2):

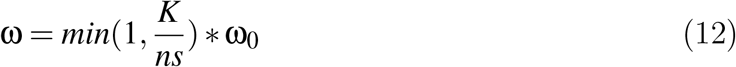

We further modified the policy to parameterize additional processes. For instance, individuals often make value-independent, random lapses in choice while doing the task. To capture this property of behavior, we derived a secondary policy by adding a random noise parameter in choice selection (Nassar & Frank, 2016):

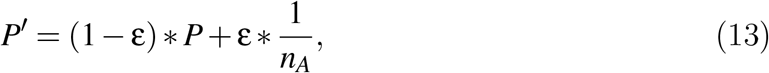

where *n*_*A*_ is the total number of possible actions, 1*/n*_*A*_ is the uniform random policy, and ε is the noise parameter capturing degree of random lapses.

We fit the different configurations of the full RL-WM model to the data from Experiment 2, where we varied the set size, which permits us to modulate involvement of WM.

In the absence of a set size manipulation, it is not possible to separately identify WM from RL modules. Thus, in the first experiment, where set size is fixed, we only consider the RL module as approximating the joint contributions of both, and do not include a WM module. Because the RL module summarizes both RL and WM contributions, we add to it a short-term forgetting feature of the RL-WM’s WM module: specifically, we implemented decay in Q-values for all cards and all choices at each trial:

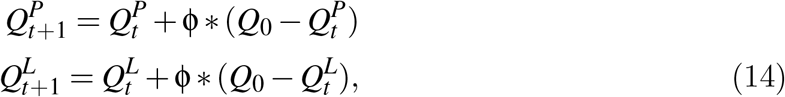

whereas in the RL-WM model the forgetting is limited to the WM module only. The list of baseline parameters for RL+WM model (Experiment 2) included: learning rate (**α**), inverse temperature (**β**), lapse (**ε**), learning bias (LB), decay (**φ**), capacity (K), WM weight (**ω**),and value mixture (**ρ**). The baseline RL model (Experiment 1) included learning rate (**α**), inverse temperature (**β**), lapse (**ε**), learning bias (LB), decay (**φ**) and value mixture (**ρ**). We explored different model variants by making different parameters fixed/varied across conditions. In RL-WM (Experiment 2) model, the parameters did not vary as a function of set size (i.e. same label/position parameter values for all set sizes).

#### 2.5.4 Model fitting and comparison

Fitting Procedure. In both Experiment 1 and Experiment 2 modeling, we used maximum likelihood estimation to fit participants’ individual parameters to their full sequence of choices. All parameters were bound between 0 and 1, with the exception of β parameter, which was fixed to 100 (found to improve parameter identifiability here and in previous similar tasks (Master et al., 2020)), and the capacity parameter (K) of Experiment 2 models, which could take on one of the discrete values ranging from 2-5. To find the best fitting parameters, we used 20 random starting points with MATLAB’s fmincon optimization function (Wilson & Collins, 2019).

##### Model validation

To validate whether our models could indeed capture the behavioral properties we set out to model, we simulated performance from the best parameter estimates for each subject 100 times per subject. We then compared whether the model predictions from the simulated data captured the patterns we observed in the actual data set.

These simulations also allowed us to ensure that our fitting procedure could adequately recover parameters in our experimental context, by fitting the model to the simulated data and evaluating the match between the true simulation parameters and recovered parameters fit on simulated data.

##### Model comparison

Exploring the full model space would lead to a combinatorial explosion of models, given the possible variations along all parameters. Thus, to explore the model space, we took a systematic approach by starting with the most complex model (all parameters varied across conditions), and gradually decreasing model complexity, while also monitoring the goodness of model fit. Specifically, we reduced the model complexity only if we found that removing a parameter improved the model fit. We chose this approach in order to conduct model comparison systematically - by testing out plausible parameter configurations with varying complexity. We compared the models using Akaike Information Criterion (AIC) (Wagenmakers & Farrell, 2004), which evaluates model fit using likelihood values and applies a complexity penalty based on the number of parameters. To ensure that our models were identifiable with AIC, we computed a confusion matrix (Wilson & Collins, 2019) by creating synthetic data sets from each model, and fitting each model to the simulated data set (computing AIC scores for each fit, see supplementary materials). This confirmed that AIC was adequately penalizing for model complexity in our situation.

## 3 Results

### 3.0.1 Experiment 1: Behavioral results

We first asked whether participants learned differently across experimental conditions. Learning curves show that participants learned well in all conditions, as their accuracy increased with more exposure to each card (Fig. 3A). A repeated measures one-way ANOVA confirmed that there was a main effect of condition (label/position/control) on performance (*F*(2, 61) = 97.7, *p* = 4.5*e −* 26). We next tested which specific conditions contributed to the significant difference, and found that the control and position conditions were not statistically distinguishable (paired t-test: *t*(61) = 1.61, *p* = .11). Thus, the additional choice feature (the labels) in the position condition did not seem to affect the choice process. The label condition performance, however, was significantly lower than the position and the control condition performances (paired t-test: position: *t*(61) = 11.1, *p* = 3.8*e−* 16; control: *t*(61) = 12.9, *p* = 5.4*e−* 19).

**Figure 3:**
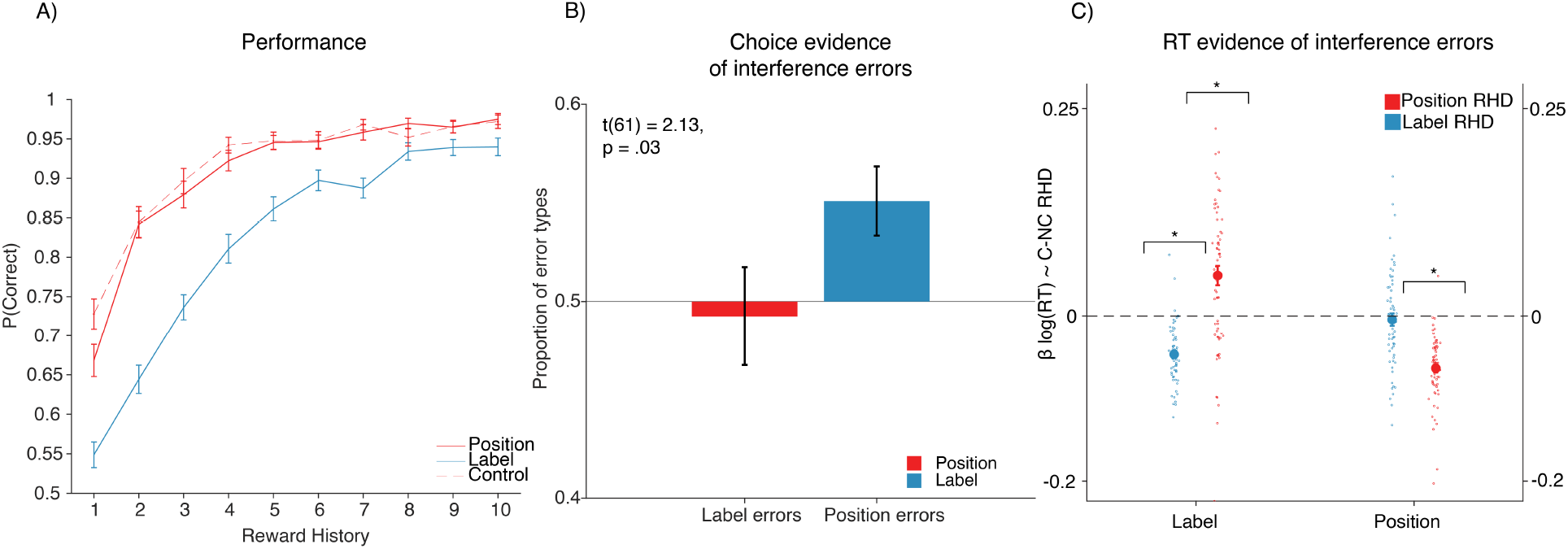
Experiment 1 Model-Free results. A) Proportion of correct choices as a function of number of previous rewards obtained for a given stimulus. Participants performed worse in the label condition, compared to the position and control conditions. Performance in the position and control conditions did not differ statistically. B) Asymmetric value interference: The values of motor actions interfered with values of correct labels in the label condition, thus resulting in the interference errors, but not the other way around. C) Mixed effects regression model shows that the interference of motor action reward history/values may have resulted in the delayed response/longer RTs in the label condition. * indicates statistical significance at *p <* .05

We next examined why label condition performance was worse. We hypothesized that choice was not simply noisier in the label condition, but instead that choice might be contaminated by the reward history of irrelevant motor choices. To test this hypothesis, we computed the cumulative card-dependent label/position reward history (see methods), and quantified the proportion of error trials in which participants incorrectly chose a box with high reward history of an incorrect feature (Fig. 1B). In the position condition, participants did not make more interference errors than expected by chance level (0.5 for two possible errors) (Fig. 3B; *t*(61) = .13, *p* = .89). This confirms that the presence of labels in the position condition did not impact choice compared to the control condition. By contrast, in the label condition, the proportion of interference errors was significantly higher than chance (Fig. 3B; *t*(61) = 2.54, *p* = .01). Furthermore, the proportion of interference errors in the label condition was significantly greater than interference errors in position condition (*t*(61) = 2.13, *p* = .03). This result suggests an asymmetry in interference between different choice spaces, in that the values of less general/motor action choices seem to contaminate the more general choice process (but not the other way around). To rule out the possibility that the effect we observe is driven by the block/condition order (i.e. transfer of incorrect strategy from the previous block), we ran a mixed-effects general linear model predicting accuracy with previous vs. current block conditions. The result of this analysis showed that participants’ performance was affected by the current block condition (*p* = 2.22*e−* 14), but not the previous block condition (*p* = 0.45), thus ruling out order effects as a possible explanation of our results.

Next, we performed a trial-by-trial analysis to examine the effect of card/label values on correct trials’ reaction times (*RT*). For each condition, we used a mixed effects linear model to predict log(*RT*) from the reward history difference (RHD) between chosen and unchosen choices (see methods), where choice referred to label in one predictor and position in the other. The rationale behind this analysis is that if participants are engaging the appropriate decision strategy, then RTs should decrease with the higher RHD in the condition-relevant dimension (label or position), because higher RHD means greater evidence in favor of the correct response. On the other hand, in the event of interference, we expected participants’ RTs to be modulated by the RHD of the incorrect dimension (e.g. position RHD in label condition). We controlled for the trial number in the model.

As predicted, participants’ RTs decreased with increased respective RHD in both label (Fig. 3C; β_*label*_ = *−*.04, *p* = 5.1*e−* 19), and position conditions (Fig. 3C; β_*position*_ = *−*.06, *p* = 3.6*e−*21). Label RHD did not affect the RTs in the position condition (β_*label*_ = *−*.004, *p* = 0.55). Hence, the mixed effects model aligned with interference errors, confirming that participants’ choices were not affected by the presence of additional feature (the labels) in the position condition. On the other hand, the position RHD surprisingly increased RTs in the label condition (β_*position*_ = .034, *p* = .001), suggesting that the interference of motor action values with label values may have resulted in the delay of choices (Fig. 3C). We compared the subject-level β estimates of the effect of incorrect dimension RHD on RTs in position and label conditions, and found that the incorrect RHD effect was significantly greater in the label condition (paired t-test: *t*(61) = 3.87, *p* = 2.6*e−* 04), confirming the asymmetry between conditions revealed in previous analyses.

### 3.0.2 Experiment 1: Modeling results

We used computational modeling to tease apart the mechanisms driving the condition effects. We fit several variants of reinforcement learning (RL) models, and focus here on 4 models that represent the main different theoretical predictions (Fig. 4A; Fig. 4B). The standard RL model (M1) assumes no difference between the conditions and serves as a baseline that cannot capture the empirical effect of condition. RL model M2 lets learning rates depend on condition, and tests the prediction that slower learning with labels is driven by different rates of reward integration. Model M3 extends model M2 with an additional mechanism, parameterized by the value mixture (ρ_*L*_), that enables position value to influence policy in the label condition.

**Figure 4:**
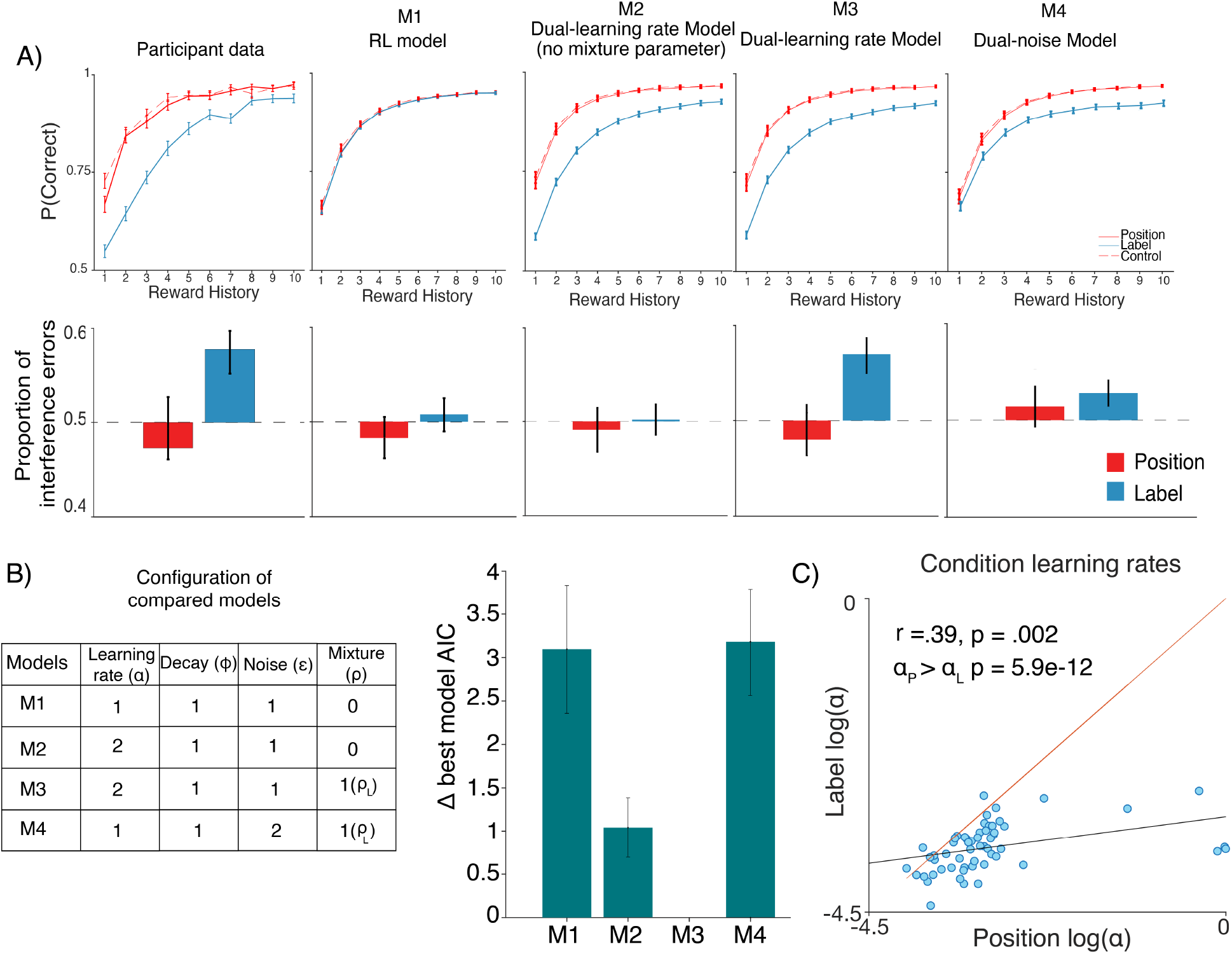
Experiment 1: Modeling Results. A) Model validation comparing the observed data to predictions of tested models; M3 reproduces behavior best. B) Parameters used in models M1-4 (left); M3 has best group-average AIC. C) Comparison of condition-dependent learning rates shows that learning rates are correlated, and that label condition learning rates are significantly lower compared to position condition learning rates.

Ruling out difficulty explanation using computational modeling. Model M4, the dual-noise model, is an RL model with condition-dependent noise parameter (**ε**). M4 captures the hypothesis that the label condition is more difficult, resulting in a noisier choice process. Models M1-4 all assume ρ_*P*_ = 1, with no influence of labels in position blocks. Other models considered separate decay (**φ**) parameters and free position condition ρ_*P*_, but did not improve fit (see supplementary materials Fig. 7).

Model M3 offered the best quantitative fit to the data, as measured by AIC (Fig. 4B). Furthermore, only model M3 was able to qualitatively reproduce patterns of behavior. Specifically, for each of the models, we simulated synthetic data sets with fit parameters and tested whether the model predictions matched the empirical results. We focused on 2 key data features in our model validation: performance averaged over the stimulus iterations (learning curves), and asymmetrical interference errors. Model validation showed that only the model with 2 learning rates and one **ρ** parameter (M3) captured both properties of the data (Fig. 4A). These results confirm that the learned value of (irrelevant) motor actions influenced the selection of more general label choices. Further-more, model comparison results also show that slower learning in the label condition was not due to a noisier choice process, but due to reduced learning rate. Indeed, tunchanged, when controlling for WM contributionshe position condition **α** was significantly greater than label condition **α** (sign test; *p* = 5.9*e−* 12, Fig. 4C). Interestingly, the learning rates in the two conditions were correlated (Spearman **ρ** = .39,*p* = .003; Fig. 4C), suggesting that the learning process in the two conditions was driven by related underlying mechanisms.

### 3.0.3 Experiment 2: Behavioral Results

The results of the first experiment suggest that the choice type affects learning. However, given the experimental design, our conclusions could not dissociate whether the difference in RL parameters actually reflected a difference in RL mechanisms or in WM mechanisms. Recent work (A. G. Collins, 2017; A. G. E. Collins & Frank, 2018), nevertheless, suggest that RL behavior recruits other learning systems, such as WM. Hence, the variations that may appear to be driven by RL mechanisms might conceal what is actually a WM effect. To address the question of whether the choice definition matters for learning at the level of RL or WM, and whether slowed learning stems from slowed WM or RL, we ran a second experiment. In Experiment 2, we varied the number of cards (set size) to manipulate WM involvement. Furthermore, we fit variants of RL+WM model to test the contribution of WM mechanisms.

Experiment 2 results replicated findings from the first experiment, showing that there was a main effect of condition (Fig. 5A; repeated measures one-way ANOVA (*F*(1, 56) = 98.95, *p* = 5.52*e −* 14)). Furthermore, we replicated the pattern of interference errors suggesting that the value of position choices interferes with that of label choices, but not the other way around (Fig. 5B; *t*(55) = 2.89; *p* = .006).

**Figure 5:**
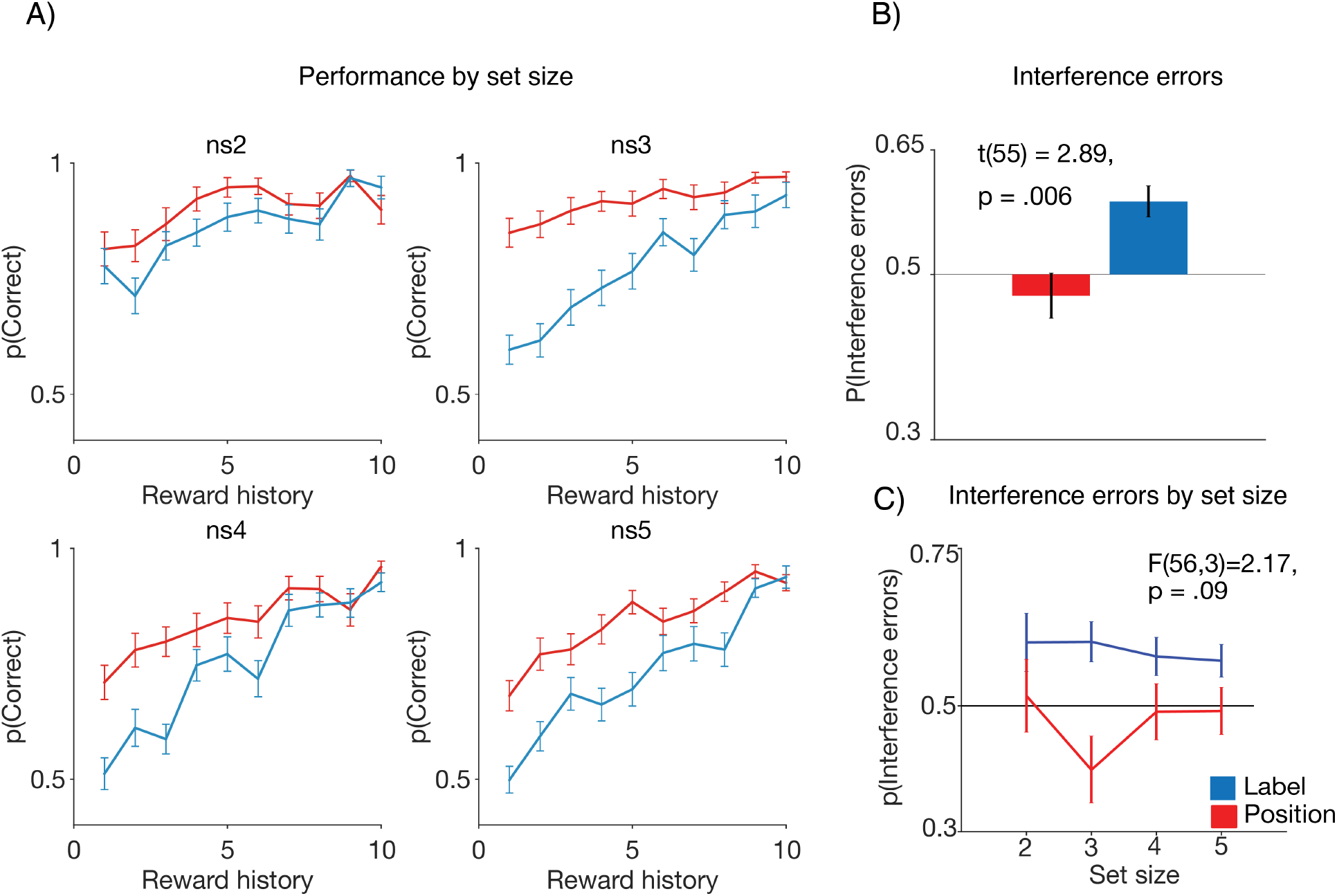
Experiment 2 Results. A) Participants’ overall performance varied by set size (a marker of WM contribution), and was worse in the label condition. B) The asymmetry in value interference replicated from first experiment, showing that values of position choices interfere with values of label choices, but not the opposite. C) The interference errors did not vary by set size.

We next investigated how set size manipulation affected these results. As predicted, performance decreased with set size in both conditions (Position: *F*(3, 56) = 11.83, *p* = 4.51*e−* 07; Label: *F*(3, 56) = 23.498, *p* = 9.45*e−* 13). There was an interaction between set size and condition (*F*(3, 56) = 16.21, *p* = 5.92*e−* 10; Fig. 5A). There was no set size effect in interference errors (*F*(3, 56) = 2.17, *p* = .09, Fig. 5C).

To better understand the source of the set size effect, we ran a general linear mixed effects model to predict trial-by-trial performance. As with the mixed effects model in Experiment 1 data analysis. Our mixed-effects model included predictors indexing WM mechanisms (set size and delay between current stimulus and the most recent rewarded stimulus presentation; indexing capacity and susceptibility to decay properties of WM respectively), and RL effects (dimension-relevant, card-dependent reward history, calculated from the cumulative number of earned points for each card, indexing reward-based learning). We also ran a model which tests for an interaction between individual RL/WM factors and the task condition.

A likelihood ratio test provided evidence in favor of the interaction model over a model without interactions (*p <* .05). The interaction model showed that, as expected, participants’ performance increased as a function of reward history (β = .62, *p* = 9.7*e−* 30), and decreased as a function of set size (β = *−*.18, *p* = .00011). There was no effect of block (β = .04, *p* = .58) or delay (β = *−*.04, *p* = .37), suggesting that neither overall task exposure nor delay, affected performance over and above reward history and set size. The only significant interaction term was the condition*reward history interaction (β = .16, *p* = .01), suggesting that the reward history more heavily contributed to an increase in performance in the label condition. To understand our results on a more mechanistic level, we turn to computational modeling.

### 3.0.4 Experiment 2: Modeling results

The set size manipulation in experiment 2 enables us to identify distinct contributions of RL and WM (A. G. E. Collins & Frank, 2012) with the full RL-WM model (see methods). Briefly, RL-WM disentangles an incremental, value-learning process (RL), as well as a rapid-learning, but decay-sensitive short-term memory-based decision process (WM). Choice policy is a weighted mixture of RL and WM (Fig. 2A,B), where the weighting is proportional to one’s WM capacity. In other words, the model architecture posits that if one’s WM capacity is low, one might be more likely to rely on RL than WM, especially when set size (number of items) is high. We first replicated in experiment 2 that models including only a single of those mechanisms cannot capture the set size effect adequately, as has been shown before (A. G. E. Collins & Frank, 2012). We then approached model comparison by systematically varying the complexity of the RL+WM model (Fig. 2A), in order to establish whether specificity in RL or WM module parameters (or both) is necessary to capture the divergence between behavioral patterns in 2 conditions. Because the RL-WM model assumes policy for choice generation at the level of both RL and WM, we also tested if integrating irrelevant dimension interference with a **ρ** mixture parameter in the policy of RL module or WM module (or both) captures our data the best. We were interested in the condition-based dissociation between parameters.

Exploring all possible parameter combinations was computationally prohibitive. Thus, we explored a subset of the most relevant models (see methods; in the main text, we focus only on a subset of models; see supplement for other non-winning models - Fig. 9). Using AIC comparison, we identified the simplest model which allowed us to capture the properties of the data (M1, (Fig. 6A)). In M1, the WM weight (**ω**) and **ρ** parameters were condition-dependent (with free **ρ** parameter for label condition, and position condition **ρ** fixed to 1). Capacity (K), learning rate (α), decay (φ), learning bias (LB), noise (ε) were shared across the 2 conditions - model comparison showed no benefits to making them independent (Fig. 9). We further consider 3 other variants of this model: no value interference **ρ** (M2), **ρ** in RL policy alone (M3), and **ρ** in WM policy alone (M4) (Fig. 6A). Last, we consider a control model with condition-dependent ε and α, which would attribute the decline in label performance to noise/RL system primarily (M5). Consistent with Experiment 1 results, the AIC comparison revealed that M5 could not capture data well, and that M1 without **ρ** (M2) fit worse (Fig. 6A), providing additional evidence for the necessity of the interference mechanism to capture choice data, and thus, the existence of motor value interference in label blocks. However, the AIC comparison failed to significantly distinguish between the remaining models M1 (ρ in RLWM), M3 (ρ in RL) and M4 (ρ in WM) (repeated measures ANOVA: *F*(2, 56) = 2.63,*p* = .07), though **ρ** in RL models fit numerically worse, supporting the idea that we needed to include motor value interference in the WM module to account for the results. Therefore, we henceforth focus on the simplest model, M1 with condition-dependent **ω** and **ρ** in RL and WM policy, as this model makes the fewest specific assumptions about RL-WM dissociation between the 2 conditions. Note that model comparison results were identical (and stronger) when using BIC instead of AIC, and that protected exceedance probability supported M1 over other models (see supplement Table 1 and Table 2, Fig. 14).

**Table 1.** Protected Exceedance Probability of tested models in Experiment 2, computed based on AIC evidence. Bayes Omnibus Risk *BOR* (indexing the probability that model frequencies are equal) = 0.94, which suggests that frequency is not strongly differentiable between models.

| M1 | M2 | M3 | M4 | M5 |
| --- | --- | --- | --- | --- |
| 0.20 | 0.18 | 0.19 | 0.22 | 0.18 |

**Table 2.** Since BIC provided stronger differentiation between models, we computed the protected exceedance probability based on BIC evidence. Bayes Omnibus Risk (*BOR*) = 1.29*e−* 12, with *PXP*(*M*1) = 1,suggests that M1 has the highest frequency.

| M1 | M2 | M3 | M4 | M5 |
| --- | --- | --- | --- | --- |
| 1 | 0 | 0 | 0 | 0 |

**Figure 6:**
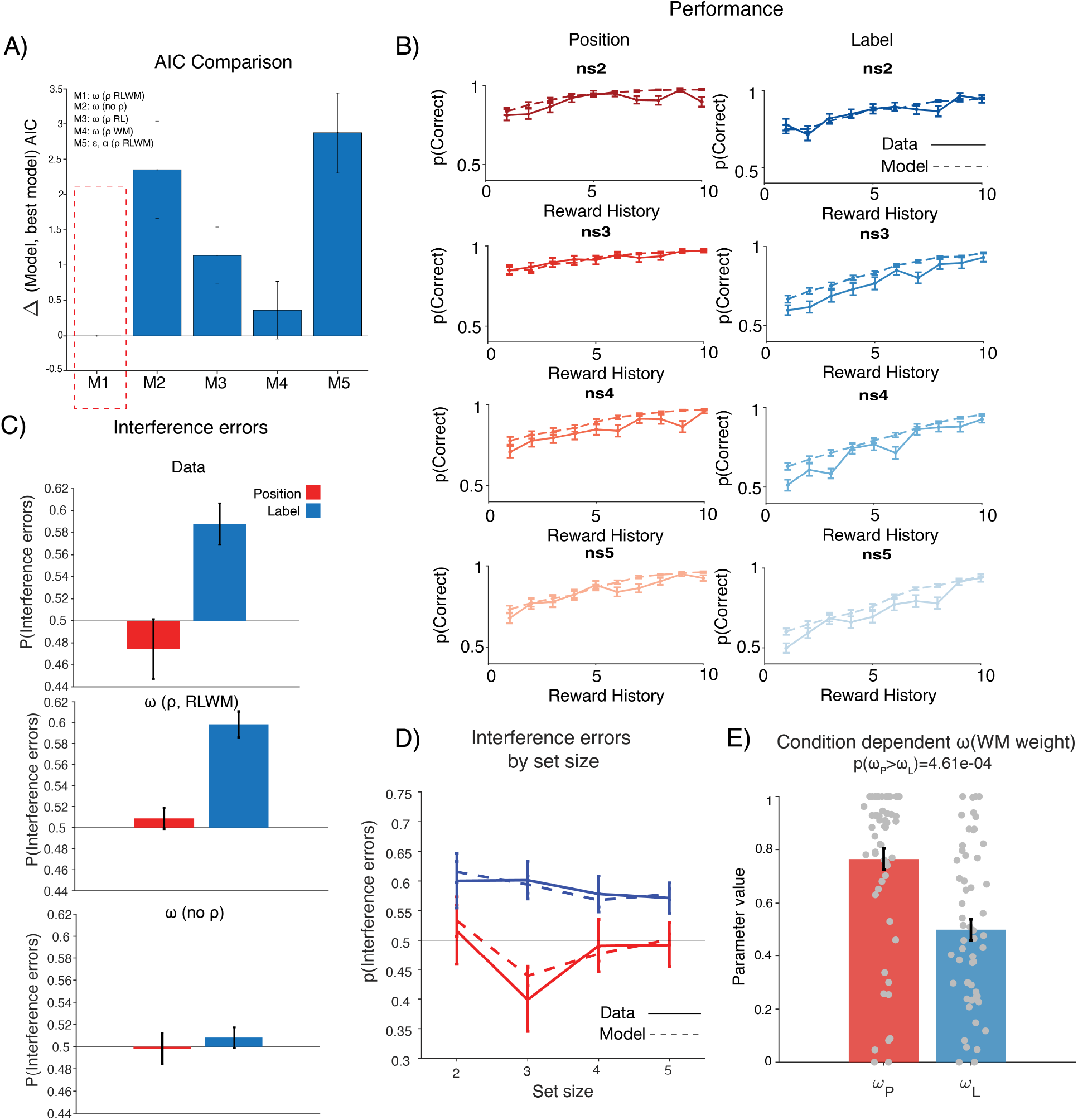
Experiment 2: Modeling Results. A) AIC comparison allowed us to narrow down the space of models. Models with condition-specific WM weight (**ω**) fit the best (M1-M4). Removing the mixture parameter (ρ) harmed the model fit (M2). A model assuming impairment in RL did not fit as well (M5). See main text for model specifications. B) Model simulations of the best model M1 captured the behavioral data patterns. C) Model validation for M1 (ρ) and M2 (no ρ) confirms the necessity of **ρ** parameter in capturing the interference error patterns. D) M1 captured interference errors in different set sizes. E) Comparison of condition-dependent parameters shows that **ω** is lower in the label condition.

The M1 model adequately captured the data patterns in 1) learning curves (Fig. 6B), 2) overall interference errors (Fig. 6C) and 3) interference errors by set size (Fig. 6D). Furthermore, the WM weight **ω** was significantly reduced in the label condition compared to the position condition in M1 (Fig. 6E).

Overall, the results suggested that the performance decrease in the label condition was driven primarily by deficits in WM, specifically by a smaller WM weight **ω** that indexes the set-size independent contribution of WM to learning. Therefore, the choice type (more/less general) impacted learning, and it seemed to do so by decreasing participants’ ability to use WM for learning. However, the value interference appeared to be present in both RL and WM mechanisms.

## 4 Discussion

Humans and animals make many types of choices, at multiple levels of generality, where some choices are dependent on others. We designed a new experimental protocol to investigate whether and how different choice types impact learning. Across two experiments, behavioral analyses and computational modeling confirmed our prediction that the generality of choice type impacts learning, with worse performance for choices that do not map on to a simple motor action. Computational modeling revealed two separable sources of impairment. First, value learning for relevant choices of a more general type was slower, as revealed by smaller learning rates (α) in experiment 1. Second, choices were contaminated by irrelevant motor action values. The second experiment examined whether this dissociation originated in different neuro-cognitive systems’ contributions to learning, namely RL and/or WM. Our results revealed that the reduction in learning speed for general-format choices stemmed more from WM than the RL process, with WM weight (**ω**) reduced but RL (α) unchanged, when controlling for WM contributions. However, the interference of low level value appeared to be present in both mechanisms. The selective reduction in WM weight implies that participants’ executive resources might be leveraged to define the choice space that is then used by both the RL and WM system; a more generalized choice space requires a higher degree of such computation, thus leaving reduced resources for actual learning.

In both experiments, we found an asymmetry in interference between choice types. When participants learned to make more general choices (selecting a label) that required a subsequent motor action (pressing the key corresponding to the label’s location), their choices were influenced by the irrelevant reward history of motor actions. By contrast, when participants learned to make less general choices (correct response is defined by pressing the same key corresponding to the box location), they were not influenced by the irrelevant reward history of label boxes. This result is consistent with a choice hierarchy interpretation, where participants may be unable to turn off credit assignment to irrelevant choice dimensions when the realization of their (abstract) choice does involve this dimension (Eckstein & Collins, 2020), but are able to do so when the irrelevant choice dimensions are more abstract, as shown here.

While our evidence implies that participants exhibit a decision bias towards motor actions, we acknowledge that our protocol cannot disambiguate between the motor actions themselves and the corresponding spatial location of the boxes. That is, we cannot confirm whether the participants track the value of specific motor actions (index/middle/ring finger key press) or of the corresponding box positions (left/middle/right). Hence, a competing interpretation of our results would be that spatial position, rather than motor actions, are prioritized in tracking value, compared to other visual features such as labels. To completely rule out this possibility, we would need to modify the current task with a condition where the motor actions are not aligned with the specific positions, and inspect whether the interference effect persists in such condition. However, we think this account is less likely than a choice abstraction account, which explains our results more parsimoniously, without requiring a “special status” for a “position” visual feature.

Furthermore, animal research supports this interpretation, as it shows differences in the neural code of choices, which are defined primarily as motor actions versus more abstract choices (Rothenhoefer et al., 2017; Luk & Wallis, 2013). Specifically, these studies have utilized recordings from neurons of animals trained to perform a task that contrasted motor-action choices with stimulus goal choices, in order to identify the neural substrates which differentiate between the two. The results seem to implicate prefrontal cortex (PFC), anterior cingulate cortex (ACC) orbitofrontal cortex (OFC) and striatal regions (ventral striatum) as areas that differentiate between how choices with different levels of abstraction are coded in the brain. Therefore, it is likely that it truly is a dissociation between motor actions, rather than positions, and more abstract choices that led to the interference and the effects we observed in our work. Our results have implications relevant for the discussion on hierarchical representations. Specifically, while simple RL algorithms are useful in capturing reward-based learning they are commonly criticized since they fail to capture robustness of human learning. Hierarchical reinforcement learning (HRL) was developed in part to address these pitfalls of RL (Botvinick, Niv, & Barto, 2009; A. G. E. Collins & Frank, 2013; Stolle & Precup, 2002; Xia & Collins, 2020). Previous research suggests that the choice space might be hierarchically represented, with lower level of hierarchy consisting of primitive actions, and higher level consisting of temporally extended actions (state-dependent, extended policies), also known as options (Stolle & Precup, 2002). Evidence from this research suggests that hierarchical representations are useful for enabling transfer; instead of learning from scratch in the novel context, an agent can leverage higher level representations to speed up learning. The transfer results also suggest that choices at different levels of hierarchy show an asymmetry in flexibility in novel contexts (lower level choices being less flexible). Our results are consistent with this finding since motor actions seem less flexible and less impacted by competing reward information, providing additional supporting claims for hierarchical representations in choice space.

In addition to this, there is evidence of hierarchical representations at the neural level. In particular, frontal areas (primarily PFC) and basal ganglia (BG), are also frequently investigated as neural mechanisms which support hierarchical reasoning/learning (A. G. E. Collins & Frank, 2013). Converging insights suggest that the cortico-BG loops support representations of both low-level associations and abstract rules/task sets, giving rise to latent representations that can be used to accelerate learning in novel sittings (A. G. E. Collins & Frank, 2013; Stolle & Precup, 2002; Xia & Collins, 2020; Eckstein et al., 2019).

Both experiments implicated overall slowed learning, in addition to value interference, in the worse performance for more general choices. Our first experiment (which allowed us to test RL models only) implicated the learning rate (usually interpreted as a marker of RL system (Eckstein et al., 2021)) as the mechanism driving the difference between conditions with different choice types. However, our second experiment enabled us to test the more holistic hybrid model of RL and WM, and revealed that the impairment in the more general choice condition likely stemmed from the WM system, rather than RL. Previous work has shown that executive function (EF), in its different forms, contributes to RL computations (A. G. Collins, 2017; Niv, 2019). The general summary of this work is that high-dimensional environments/tasks pose difficulty to RL; EF then acts as an information compressor, making the information processing more efficient for RL (Rmus et al., 2020). Operating in a more generalized choice space might more heavily rely the contribution of EF (in this case WM) relative to the less abstract condition. Therefore, resource-limited WM might be leveraged to define the choice space (i.e. relevant features of the choice space-labels in label condition). As a result, the WM weight included in the WM + RL hybrid model, which indexes the WM contribution to learning, appear reduced in the label condition. Our interpretation of this result is that this reduction in WM contribution might indicate that some of participants’ limited WM resources are recruited elsewhere, and specifically that it has already been used to define the choice space over which learning and decision making occurs.

While we conclude that WM is used for defining the choice space, consistent with prior results on EF contributions to RL computations (Todd, Niv, & Cohen, n.d.), we do not make any particular assumptions about how the use of choice space is divided between RL and WM once it’s defined. We tested different model variations, with the parameter that mixes label/position values, to explain value interference at the policy level of RL, WM or both. If there was clear evidence in favor of the mixture parameter in RL or WM policy, it would imply that the policy generation based on choice space is primarily driven by that system. However, our model comparison revealed no evidence that the mixture parameter is specific to either RL or WM, suggesting that the choice space is shared between the two. This will be important to explore in future research.

A competing interpretation for our findings of slowed learning for more abstract choices is that the label condition required more attention and was more difficult. While this is true, we took steps to mitigate this potential confound on two levels - task design and modeling. In the task design, we constructed the single trial structure such that participants had a chance to see box labels first, before the onset of the card. By doing this we aimed to eliminate potential advantages of the position condition, where participants do not need to perform an additional process of identifying the label location prior to executing the response. Furthermore, our modeling enabled us to validate the effects of our task design. Specifically, in both experiments we tested the model with condition-dependent noise parameters, which predicts that different noise/difficulty levels are what drive the performance difference in our conditions. This model did not fit the data well (Experiment 1: Best model AIC > 2 noise model AIC t(56)= -5.179, p= 3.13e-06; Experiment 2: Best model AIC > 2 noise model AIC t(56)= -5.05, p= 4.98e-06), making it unlikely that difficulty-induced lack of attention/motivation could explain our condition effect.

Surprisingly, we found that participants’ response times (RT) on correct trials increased as a function of position reward history difference (RHD) in the label condition. This implies that when both label and position sorting rules were in agreement as to the best choice to make (i.e. the blue box was the correct box, and was in the position that had been most rewarded so far), response times tend to be longer (the corresponding effect was not observed in the position condition, where label RHD had no effect on RTs). This is, therefore, a counterintuitive effect, as we would expect the congruent information to accelerate response execution, rather than slow it, as observed here. One possibility might be that participants do engage in a form of arbitration between selection of different response types. Specifically, they might be biased to execute the motor action based on the reward history difference, as it seems to present itself as a default option based on our results. However, because they are informed that the response based on label selection is correct for the given block, they might delay the response execution, in order to override the default. Nevertheless, this is a speculation - careful modeling of response times is required to further explain this effect, which is beyond the scope of this paper. This account would also predict the highest degree of conflict in this congruent situation, rather than in situations where both rules disagree. It will be an important question to solve in future research.

Our results highlight the importance of correct credit assignment, and investigation of mechanisms which might lead to errors in the credit assignment process. Our results are consistent with the previous research suggesting that motor actions might have a stronger effect on the choice selection process than is usually considered (Shahar et al., 2019). Our modeling approach allowed us to show that the mixture of Q values at the policy level is what might lead to the interference effect/incorrect credit assignment. However, as of now, we cannot conclusively say whether the mixture happens selectively at the policy level of RL, WM or both.

Identification of correct rewarding responses is a critical building block of adaptive/goal-directed behavior. Impairments in one’s ability to identify the appropriate choice space, which is then used for inference process, may consequently result in mal-adaptive/suboptimal behavioral patterns. Our interference effect results suggest that some aspects of the choice space might be incorrectly overvalued, thus resulting in choice patterns that reflect repeated erroneous selection of incorrect choice types, or an inability to utilize flexible stimulus-response mapping. These kinds of perseverative responses are reminiscent of the inability to disengage from certain actions, observed in conditions such as obsessive-compulsive disorder (OCD) (Rosa-Alcázar et al., 2020). It would be interesting to use our task and computational modeling approach to investigate whether the mixture/interference of values at the policy level could also explain the behavior of such population.

In conclusion, our findings provide evidence that the choice type and how we define a choice have important implications for the learning process. The behavioral patterns (i.e. value interference from less abstract choices) are consistent with the premises of hierarchy in learning and behavior (i.e. lower levels in hierarchy impacting processing in higher levels), which has become an increasingly promising topic of research (Eckstein & Collins, 2019; A. G. E. Collins & Frank, 2013; Stolle & Precup, 2002). We also demonstrate additional evidence, relevant to the definition of the choice space, that EF (specifically WM) contributes to RL in reward-driven behaviors (Rmus et al., 2020), further demonstrating the complex interplay between various neuro-cognitive systems.

## 5 Acknowledgments

This work was funded by NSF2020844 to AGEC.

## 7 Supplementary materials

Experiment 1 additional model comparisons. We tested whether an additional decay parameter, an additional mixture parameter, a mixture parameter shared across the two conditions and free softmax temperature parameter improved the fit to the data. These models did not improve the fit compared to M3 (our winning model).

**Figure 7:**
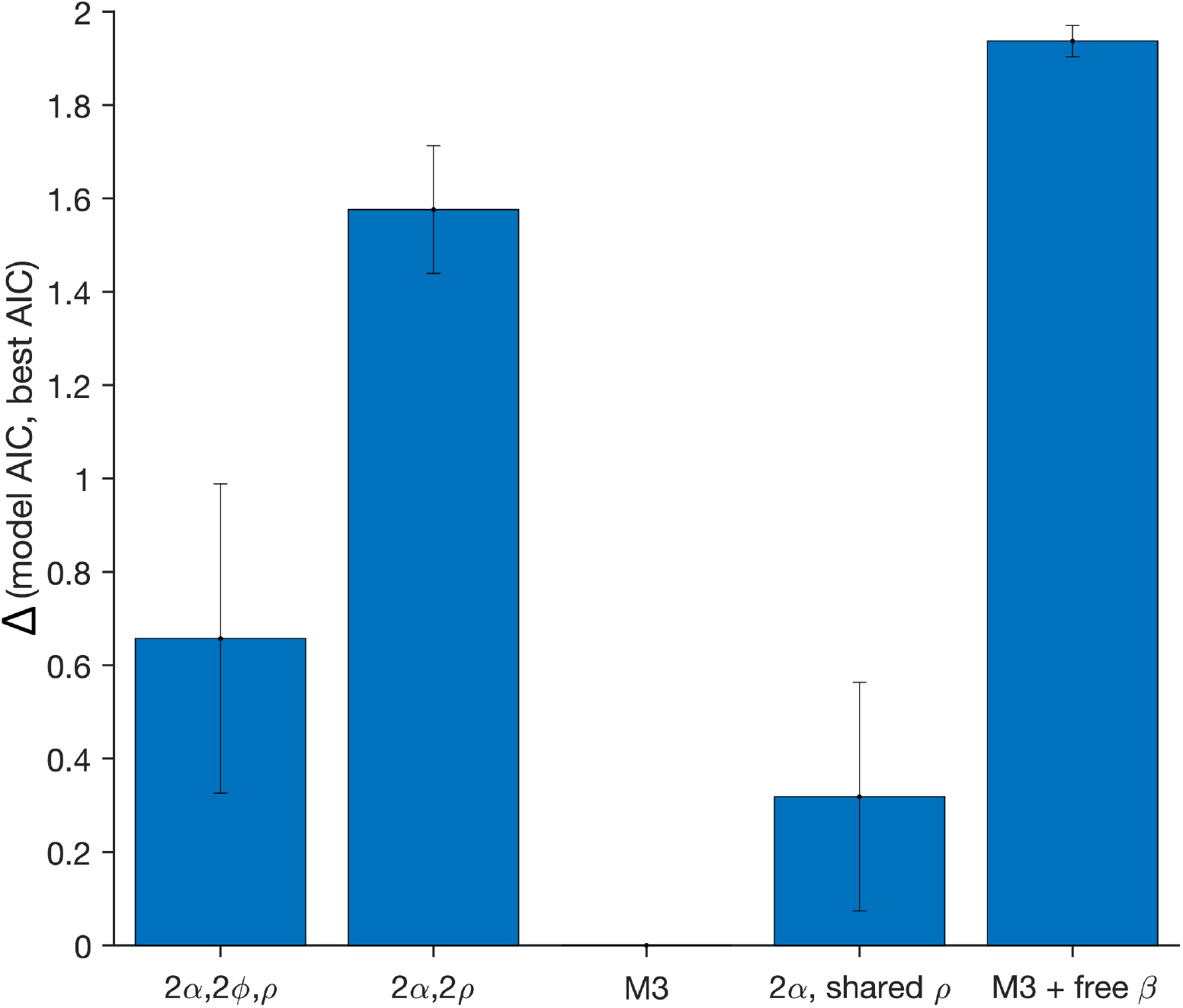
Additional models tested in Experiment 1.

Experiment 1 confusion matrix. To demonstrate the identifiability of our models (i.e. models are meaningfully different from one another), we simulated the data from each model on 62 iterations (number of participants). We used best parameter estimates for each participant to create a synthetic data set on each iteration. We then fitted each of the models to each simulated data set with 20 random starting points, to match the fitting procedure to participants’ data. Next, we computed the proportion of the times each model fit the best. If the models are identifiable, the model the data was simulated from should fit the best on most iterations (i.e. the matrix should have the highest proportion of best fit values on its diagonal). The confusion matrix showed that our models are identifiable.

**Figure 8:**
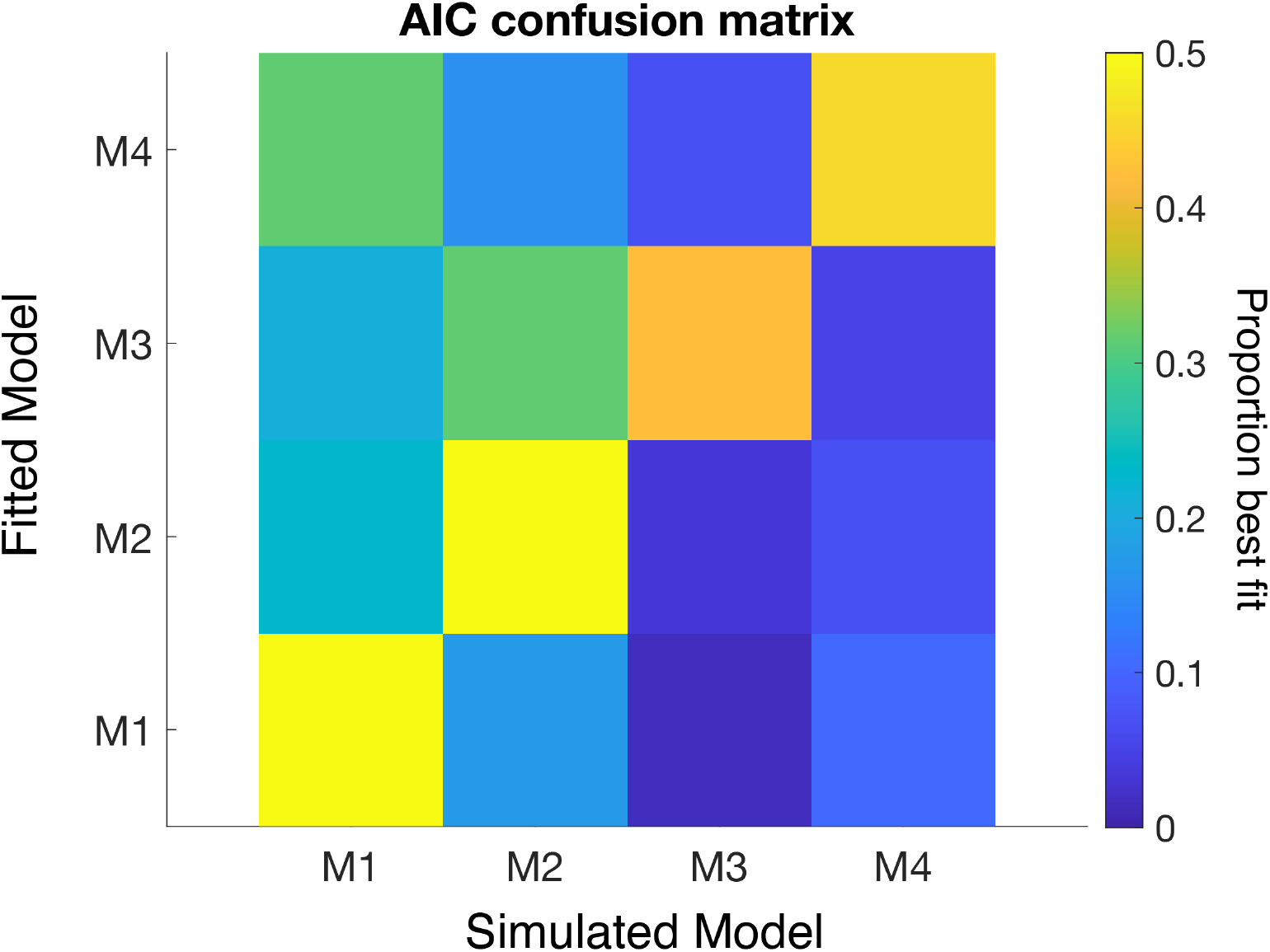
Confusion matrix of the main models tested in Experiment 1.

In our second experiment, we fit a considerable range of models, starting with the most complex (all RL + WM parameters condition-dependent), to the simplest (all parameters shared across conditions). We systematically varied the complexity of the model, while monitoring the model fit/complexity tradeoff using AIC scores, in order to test which parameters are necessary for capturing the difference between the conditions while also making sure our models are not overfitting (Fig. 9).

**Figure 9:**
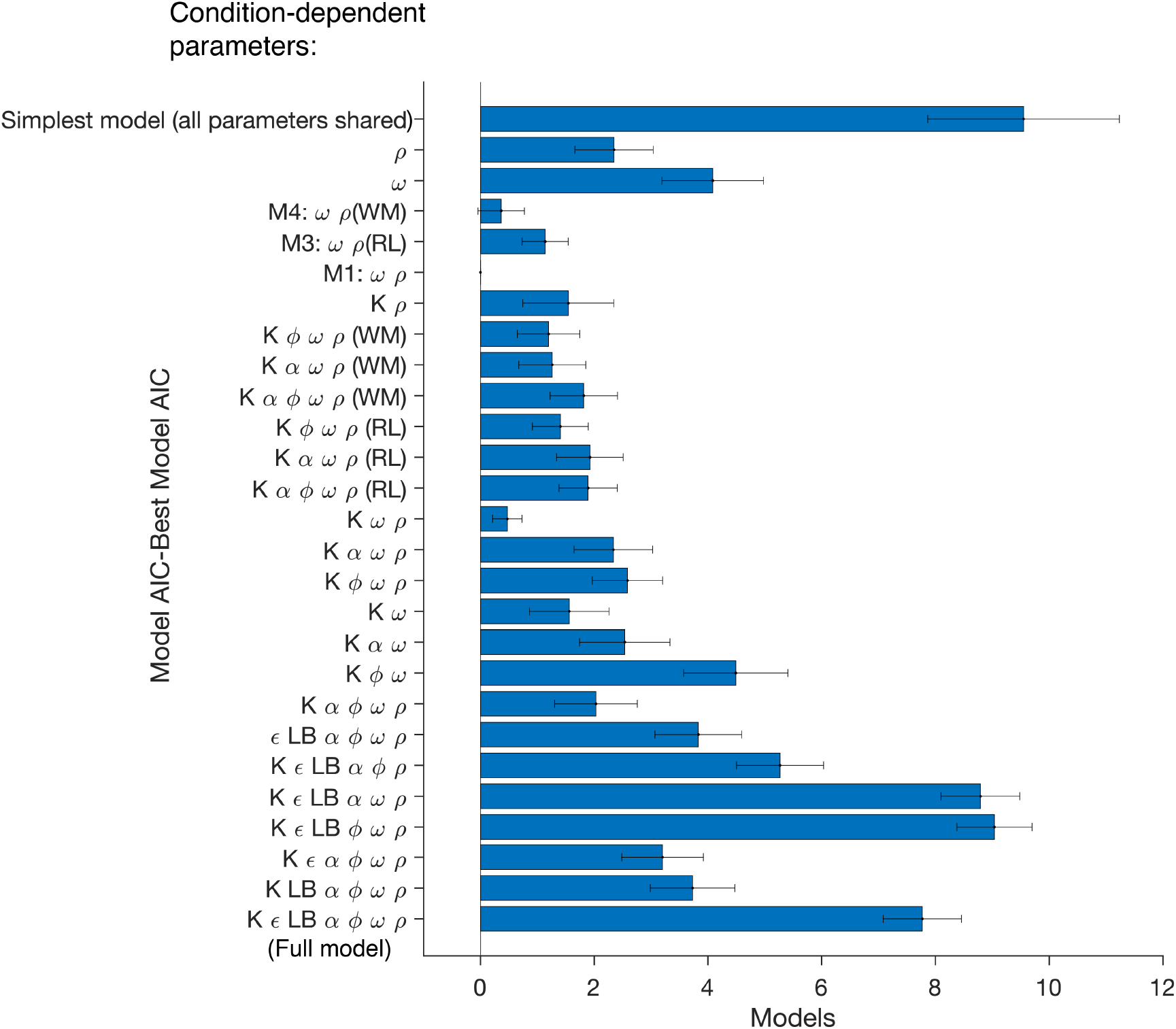
AIC comparison of models tested in Experiment 2. Here we show the difference in individual AIC scores between M3, and all other models that were tested.

Experiment 2 Confusion Matrix. We tested the identifiability of our models in experiment 2 by creating a confusion matrix (see Experiment 1 confusion matrix section in supplementary material for details on the procedure). We constructed two different confusion matrices, which test for identifiability of our model along 2 different dimensions. Our first confusion matrix allowed us to test whether the models with different placements of the **ρ** parameter (i.e. with wrong choice dimension policy mixture in RL, WM or both) are meaningfully dissociable. The confusion matrix shows that the models with mixture **ρ** in RL and WM policy can be dissociated (Fig. 10). The data simulated from the model with **ρ** parameter in both WM and RL policy was fit equally well by that model and the model with **ρ** in WM policy alone. This is consistent with our results, as model comparison revealed that AIC scores did not meaningfully differ between these two models. Note that the models included in the confusion matrix are nested models (differing by at most 1 parameter), or in the case of the second confusion matrix identical models in terms of number of parameters, but with different *rho* parameter placements. Therefore, we did not expect the AIC scores to be considerably different for paired model fits paired with data simulated across different models.

**Figure 10:**
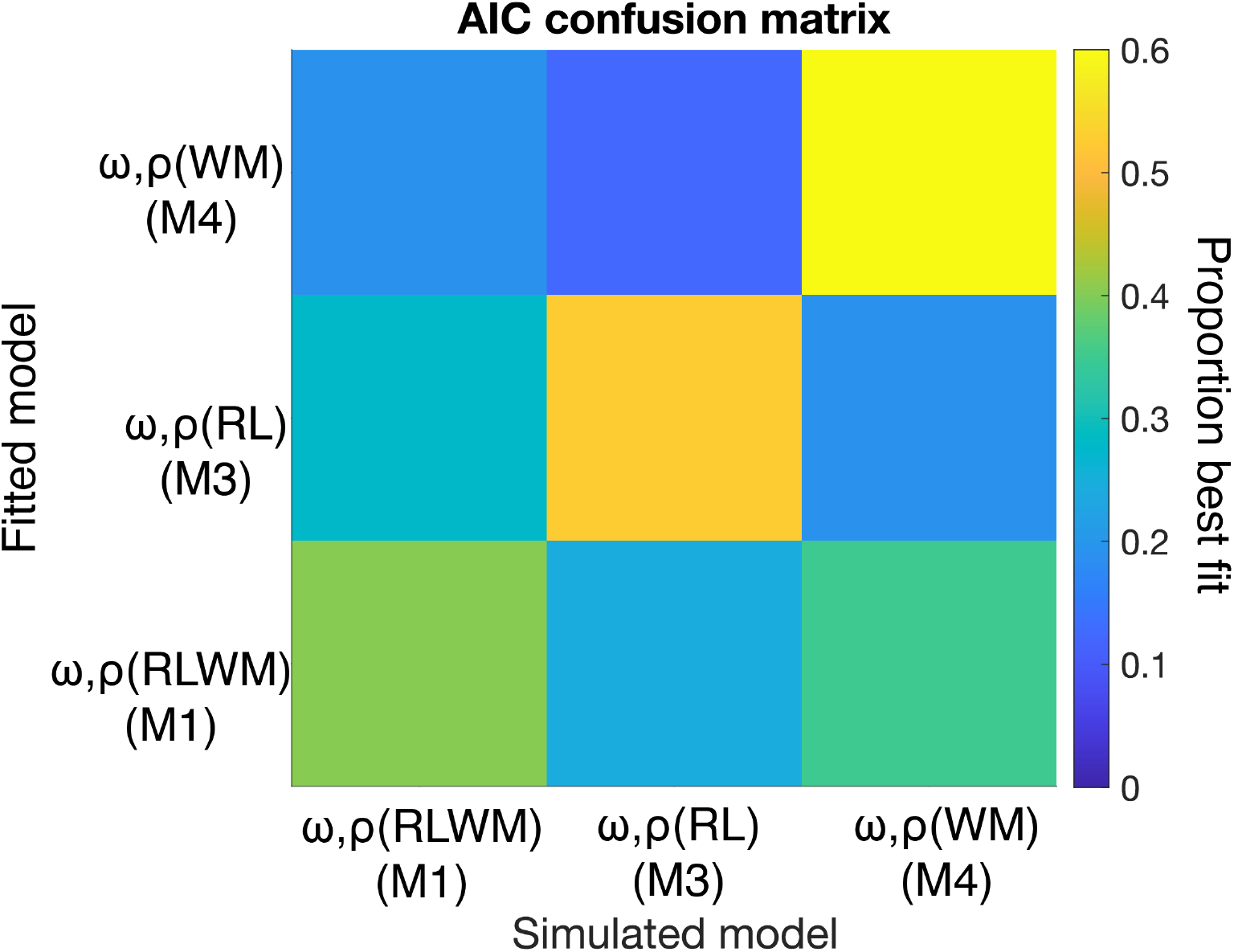
Confusion Matrix 1. Proportion of times the models fitted different simulated data sets best, based on cross-fit AIC scores for models with different placement of **ρ** paramater.

Our second confusion matrix tested whether we can dissociate the model we converged on in the main text (M1, **ω** with RL+WM ρ) from variations of model with 1) no **ρ** parameter, and 2) shared WM weight **ω**. Our results showed that our models are mostly identifiable, with an exception of M2 (Fig. 11). However, M2 cannot produce the observed qualitative error patterns, providing another method to rule out this model.

**Figure 11:**
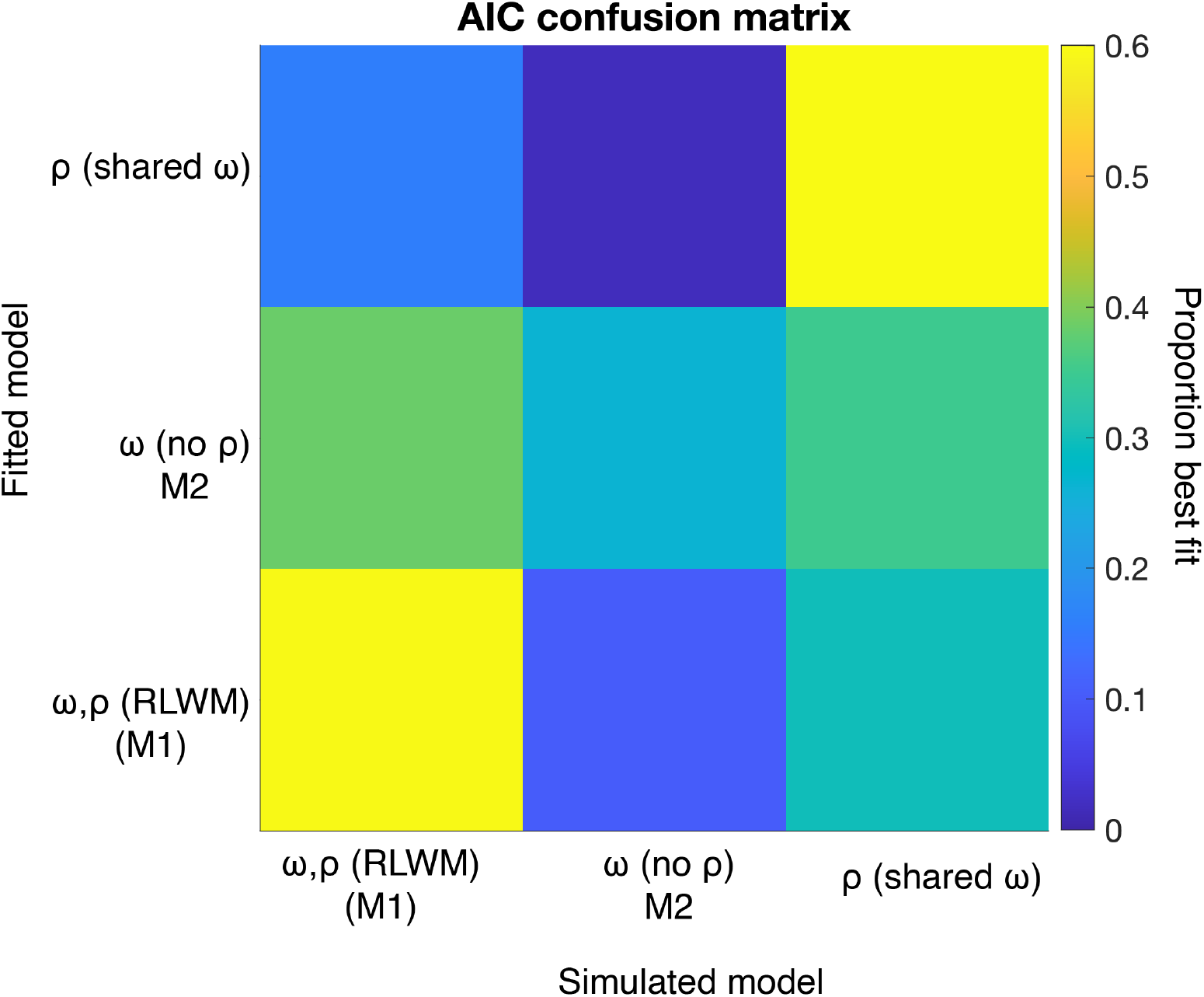
Confusion Matrix 2. Proportion of times the models fitted different simulated data sets best, based on cross-fit AIC scores for models with condition dependent **ρ** and **ω** parameters (M1), condition dependent **ω** (M2), and condition dependent ρ.

**Figure 12:**
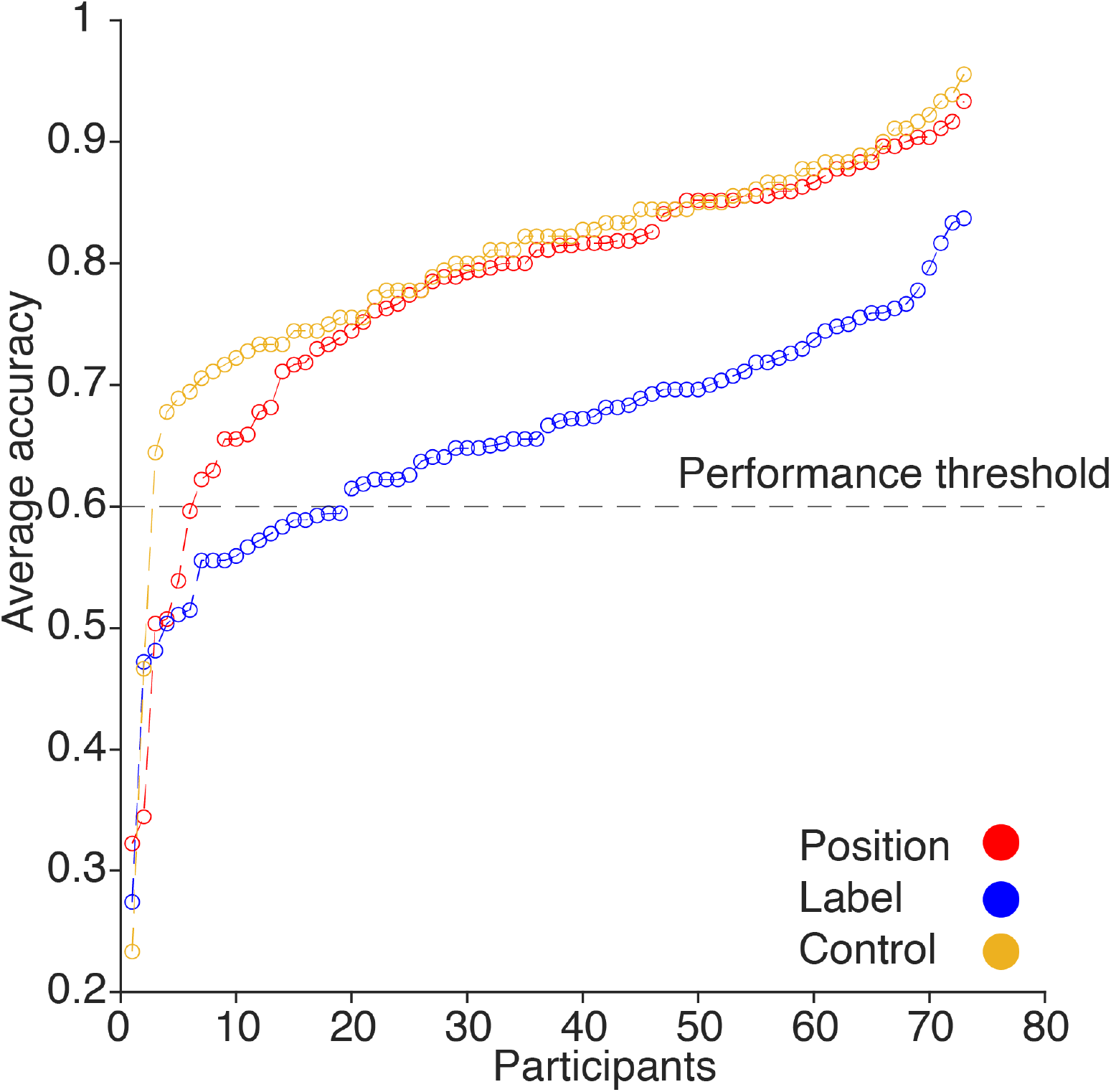
Exclusion criteria based on the task performance. We averaged accuracy across all conditions. Based on the “elbow point”, most participants’ performance is above .60, therefore we used .60 as criteria for exclusion.

**Figure 13:**
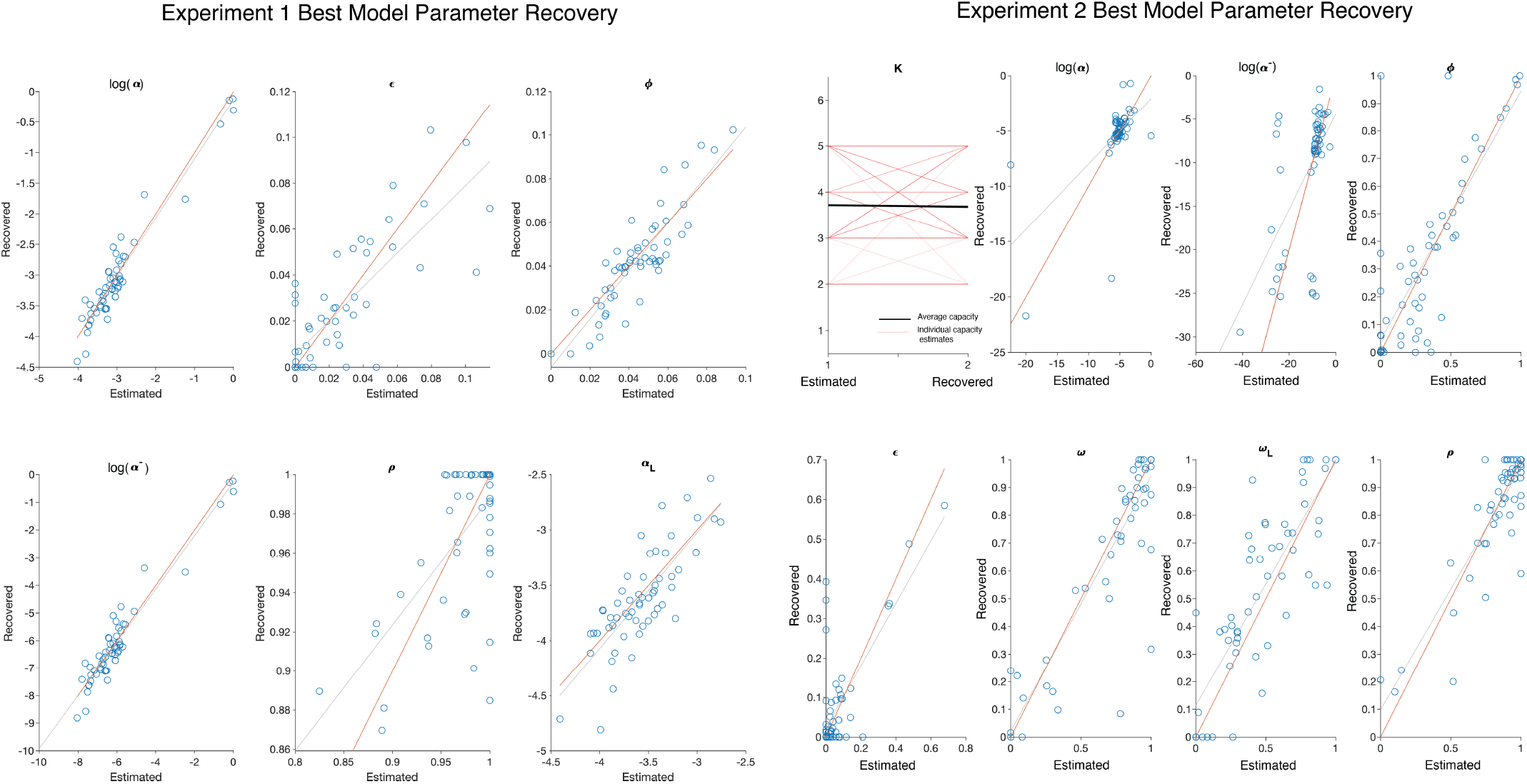
Parameter recovery for the best models in Experiment 1 and Experiment 2.

**Figure 14:**
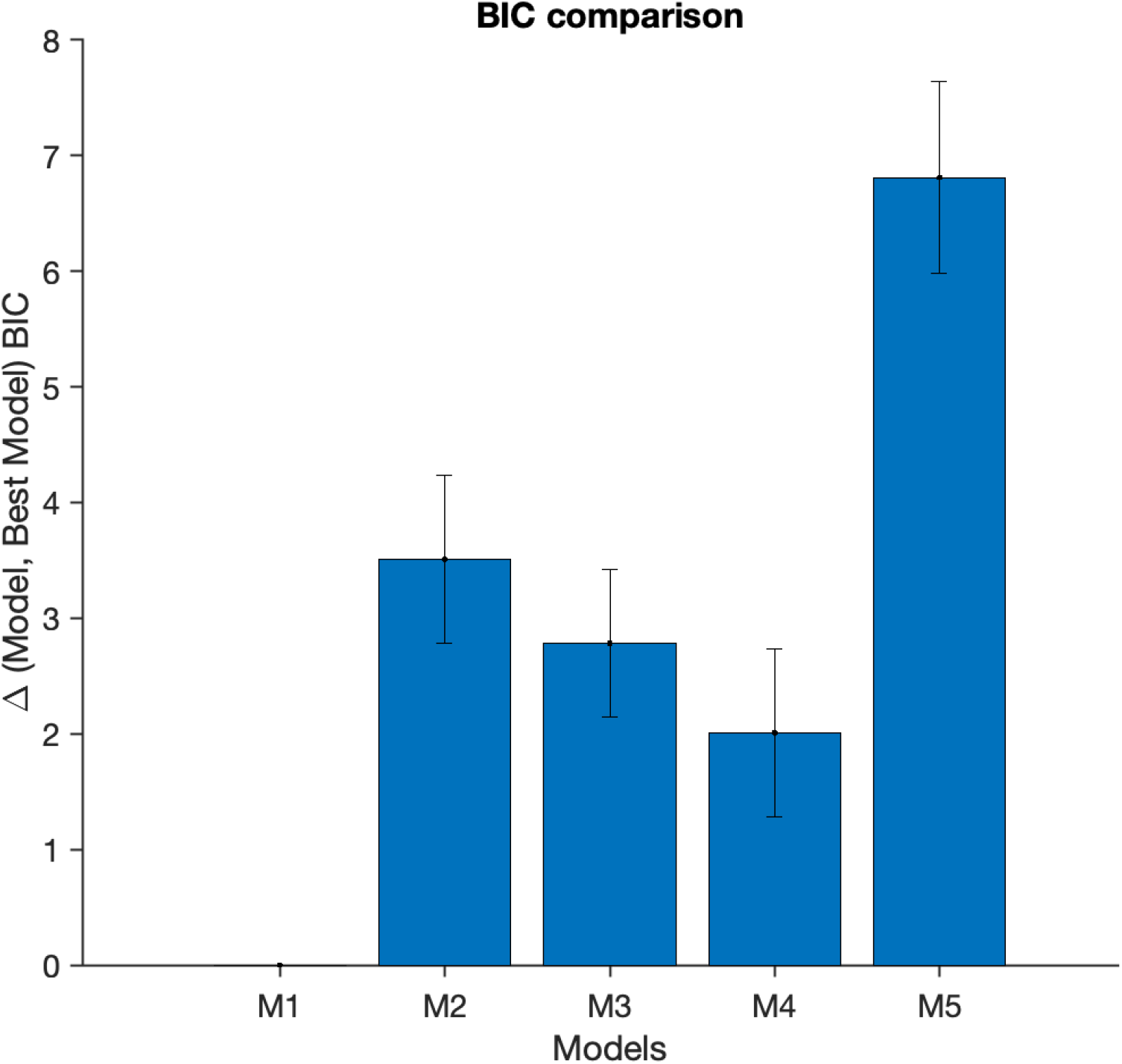
Parameter recovery for the best models in Experiment 1 and Experiment 2.

## Notes

### Competing Interest Statement

The authors have declared no competing interest.

## References

Ballard, I., Miller, E. M., Piantadosi, S. T., Goodman, N. D., & McClure, S. M. (2018, November). Beyond Reward Prediction Errors: Human Striatum Updates Rule Values During Learning. Cerebral Cortex, 28(11), 3965–3975. Retrieved 2021-07-28, from https://doi.org/10.1093/cercor/bhx259 doi: 10.1093/cercor/bhx259

Bornstein, A. M., & Daw, N. D. (2013, December). Cortical and Hippocampal Correlates of Deliberation during Model-Based Decisions for Rewards in Humans. PLOS Computational Biology, 9(12), e1003387. Retrieved 2021-07-11, from https://journals.plos.org/ploscompbiol/article?id=10.1371/journal.pcbi.1003387 (Publisher: Public Library of Science) doi: 10.1371/journal.pcbi.1003387

Bornstein, A. M., Khaw, M. W., Shohamy, D., & Daw, N. D. (2017, December). Reminders of past choices bias decisions for reward in humans. Nature Communications, 8(1), 15958. Retrieved 2021-07-22, from http://www.nature.com/articles/ncomms15958 doi: 10.1038/ncomms15958

Botvinick, M. M., Niv, Y., & Barto, A. G. (2009, December). Hierarchically organized behavior and its neural foundations: a reinforcement learning perspective. Cognition, 113(3), 262–280. doi: 10.1016/j.cognition.2008.08.011

Collins, A. G. (2017, December). The tortoise and the hare: interactions between reinforcement learning and working memory (preprint). Neuroscience. Retrieved 2021-07-07, from http://biorxiv.org/lookup/doi/10.1101/234724 doi: 10.1101/234724

Collins, A. G. E., Ciullo, B., Frank, M. J., & Badre, D. (2017, April). Working Memory Load Strengthens Reward Prediction Errors. Journal of Neuroscience, 37(16), 4332–4342. Retrieved 2021-02-08, from https://www.jneurosci.org/content/37/16/4332 (Publisher: Society for Neuroscience Section: Research Articles) doi: 10.1523/JNEUROSCI.2700-16.2017

Collins, A. G. E., & Frank, M. J. (2012, April). How much of reinforcement learning is working memory, not reinforcement learning? A behavioral, computational, and neurogenetic analysis. The European journal of neuroscience, 35(7), 1024–1035. Retrieved 2021-12-23, from https://www.ncbi.nlm.nih.gov/pmc/articles/PMC3390186/ doi: 10.1111/j.1460-9568.2011.07980.x

Collins, A. G. E., & Frank, M. J. (2013, January). Cognitive control over learning: Creating, clustering, and generalizing task-set structure. Psychological Review, 120(1), 190–229. Retrieved 2021-02-08, from http://doi.apa.org/getdoi.cfm?doi=10.1037/a0030852 doi: 10.1037/a0030852

Collins, A. G. E., & Frank, M. J. (2018, March). Within- and across-trial dynamics of human EEG reveal cooperative interplay between reinforcement learning and working memory. Proceedings of the National Academy of Sciences, 115(10), 2502–2507. Retrieved 2021-02-08, from https://www.pnas.org/content/115/10/2502 (Publisher: National Academy of Sciences Section: Biological Sciences) doi: 10.1073/pnas.1720963115

Daw, N., Gershman, S., Seymour, B., Dayan, P., & Dolan, R. (2011, March). ModelBased Influences on Humans’ Choices and Striatal Prediction Errors. Neuron, 69(6), 1204–1215. Retrieved 2021-02-08, from https://www.sciencedirect.com/science/article/pii/S0896627311001255 doi: 10.1016/j.neuron.2011.02.027

de Leeuw, J. R. (2015, March). jsPsych: A JavaScript library for creating behavioral experiments in a Web browser. Behavior Research Methods, 47(1), 1–12. Retrieved 2021-08-25, from http://link.springer.com/10.3758/s13428-014-0458-y doi: 10.3758/s13428-014-0458-y

Eckstein, M. K., & Collins, A. G. (2019, August). Computational evidence for hierarchically-structured reinforcement learning in humans. bioRxiv, 731752. Retrieved 2021-02-08, from https://www.biorxiv.org/content/10.1101/731752v1 (Publisher: Cold Spring Harbor Laboratory Section: New Results) doi: 10.1101/731752

Eckstein, M. K., & Collins, A. G. E. (2020, November). Computational evidence for hierarchically structured reinforcement learning in humans. Proceedings of the National Academy of Sciences, 117(47), 29381–29389. Retrieved 2022-03-14, from https://www.pnas.org/doi/10.1073/pnas.1912330117 (Publisher: Proceedings of the National Academy of Sciences) doi: 10.1073/pnas.1912330117

Eckstein, M. K., Starr, A., & Bunge, S. A. (2019, April). How the inference of hierarchical rules unfolds over time. Cognition, 185, 151–162. Retrieved 2021-02-08, from https://www.sciencedirect.com/science/article/pii/S0010027719300150 doi: 10.1016/j.cognition.2019.01.009

Eckstein, M. K., Wilbrecht, L., & Collins, A. G. (2021, October). What do reinforcement learning models measure? Interpreting model parameters in cognition and neuroscience. Current Opinion in Behavioral Sciences, 41, 128–137. Retrieved 2021-07-26, from https://linkinghub.elsevier.com/retrieve/pii/S2352154621001236 doi: 10.1016/j.cobeha.2021.06.004

Farashahi, S., Rowe, K., Aslami, Z., Lee, D., & Soltani, A. (2017, November). Feature-based learning improves adaptability without compromising precision. Nature Communications, 8(1), 1768. Retrieved 2021-02-08, from https://www.nature.com/articles/s41467-017-01874-w (Number: 1 Publisher: Nature Publishing Group) doi: 10.1038/s41467-017-01874-w

Foerde, K., & Shohamy, D. (2011, September). Feedback Timing Modulates Brain Systems for Learning in Humans. Journal of Neuroscience, 31(37), 13157–13167. Retrieved 2021-07-04, from https://www.jneurosci.org/lookup/doi/10.1523/JNEUROSCI.2701-11.2011 doi: 10.1523/JNEUROSCI.2701-11.2011

Frank, M. J., Moustafa, A. A., Haughey, H. M., Curran, T., & Hutchison, K. E. (2007, October). Genetic triple dissociation reveals multiple roles for dopamine in reinforcement learning. Proceedings of the National Academy of Sciences, 104(41), 16311–16316.

Gershman, S. J. (2015, October). Do learning rates adapt to the distribution of rewards? Psychonomic Bulletin & Review, 22(5), 1320–1327. Retrieved 2021-07-07, from http://link.springer.com/10.3758/s13423-014-0790-3 doi: 10.3758/s13423-014-0790-3

Luk, C.-H., & Wallis, J. D. (2013, January). Choice Coding in Frontal Cortex during Stimulus-Guided or Action-Guided Decision-Making. Journal of Neuroscience, 33(5), 1864–1871.

Master, S. L., Eckstein, M. K., Gotlieb, N., Dahl, R., Wilbrecht, L., & Collins, A. G. E. (2020, February). Disentangling the systems contributing to changes in learning during adolescence. Developmental Cognitive Neuroscience, 41, 100732. Retrieved 2021-02-08, from https://www.sciencedirect.com/science/article/pii/S1878929319303196 doi: 10.1016/j.dcn.2019.100732

McDougle, S. D., Boggess, M. J., Crossley, M. J., Parvin, D., Ivry, R. B., & Taylor, J. A. (2016, June). Credit assignment in movement-dependent reinforcement learning. Proceedings of the National Academy of Sciences, 113(24), 6797–6802. Retrieved 2021-06-15, from https://www.pnas.org/content/113/24/6797 (Publisher: National Academy of Sciences Section: Biological Sciences) doi: 10.1073/pnas.1523669113

Nassar, M. R., & Frank, M. J. (2016, October). Taming the beast: extracting generalizable knowledge from computational models of cognition. Current opinion in behavioral sciences, 11, 49–54. Retrieved 2021-07-07, from https://www.ncbi.nlm.nih.gov/pmc/articles/PMC5001502/doi: 10.1016/j.cobeha.2016.04.003

Niv, Y. (2019, October). Learning task-state representations. Nature Neuroscience, 22(10), 1544–1553. Retrieved 2021-02-08, from https://www.nature.com/articles/s41593-019-0470-8 (Number: 10 Publisher: Nature Publishing Group) doi: 10.1038/s41593-019-0470-8

Niv, Y., Edlund, J. A., Dayan, P., & O’Doherty, J. P. (2012, January). Neural Prediction Errors Reveal a Risk-Sensitive Reinforcement-Learning Process in the Human Brain. Journal of Neuroscience, 32(2), 551–562.

Poldrack, R. A., Clark, J., Paré-Blagoev, E. J., Shohamy, D., Creso Moyano, J., Myers, C., & Gluck, M. A. (2001, November). Interactive memory systems in the human brain. Nature, 414(6863), 546–550. Retrieved 2021-07-28, from https://www.nature.com/articles/35107080 (Bandiera_abtest: a Cg_type: Nature Research Journals Number: 6863 Primary_atype: Research Publisher: Nature Publishing Group) doi: 10.1038/35107080

Rescorla, R. A., & Solomon, R. L. (1967, May). Two-process learning theory: Relationships between Pavlovian conditioning and instrumental learning. Psychological Review, 74(3), 151–182. doi: 10.1037/h0024475

Rmus, M., McDougle, S., & Collins, A. (2020, June). The Role of Executive Function in Shaping Reinforcement Learning (Tech. Rep.). PsyArXiv. Retrieved 2021-07-10, from https://psyarxiv.com/9cvw3/(type: article) doi: 10.31234/osf.io/9cvw3

Rosa-Alcázar, Olivares-Olivares, P. J., Martínez-Esparza, I. C., Parada-Navas, J. L., Rosa-Alcázar, A. I., & Olivares-Rodríguez, J. (2020). Cognitive flexibility and response inhibition in patients with Obsessive-Compulsive Disorder and Generalized Anxiety Disorder. International Journal of Clinical and Health Psychology : IJCHP, 20(1), 20–28. Retrieved 2021-07-26, from https://www.ncbi.nlm.nih.gov/pmc/articles/PMC6994753/ doi:10.1016/j.ijchp.2019.07.006

Rothenhoefer, K. M., Costa, V. D., Bartolo, R., Vicario-Feliciano, R., Murray, E. A., & Averbeck, B. B. (2017, July). Effects of Ventral Striatum Lesions on Stimulus-Based versus Action-Based Reinforcement Learning. The Journal of Neuroscience, 37(29), 6902–6914. Retrieved 2021-07-07, from https://www.ncbi.nlm.nih.gov/pmc/articles/PMC5518420/ doi:10.1523/JNEUROSCI.0631-17.2017

Shahar, N., Moran, R., Hauser, T. U., Kievit, R. A., McNamee, D., Moutoussis, M., & Dolan, R. J. (2019, August). Credit assignment to state-independent task representations and its relationship with model-based decision making. Proceedings of the National Academy of Sciences, 116(32), 15871–15876.

Stolle, M., & Precup, D. (2002). Learning Options in Reinforcement Learning. In Lecture Notes in Computer Science (pp. 212–223).

Sugawara, M., & Katahira, K. (2021, February). Dissociation between asymmetric value updating and perseverance in human reinforcement learning. Scientific Reports, 11(1), 3574. Retrieved 2021-07-07, from https://www.nature.com/articles/s41598-020-80593-7 (Bandiera_abtest: a Cc_license_type: cc_by Cg_type: Nature Research Journals Number: 1 Primary_atype: Research Publisher: Nature Publishing Group Subject_term: Cognitive neuroscience;Decision Subject_term_id: cognitive-neuroscience;decision) doi: 10.1038/s41598-020-80593-7

Sutton, R. S., & Barto, A. G. (2018). Reinforcement Learning, second edition: An Introduction. MIT Press. (Google-Books-ID: uWV0DwAAQBAJ)

Tai, L.-H., Lee, A. M., Benavidez, N., Bonci, A., & Wilbrecht, L. (2012, September). Transient stimulation of distinct subpopulations of striatal neurons mimics changes in action value. Nature Neuroscience, 15(9), 1281–1289. Retrieved 2021-07-18, from https://www.nature.com/articles/nn.3188 (Bandiera_abtest: a Cg_type: Nature Research Journals Number: 9 Primary_atype: Research Publisher: Nature Publishing Group Subject_term: Decision;Neuronal physiology Subject_term_id: decision;neuronal-physiology) doi: 10.1038/nn.3188

Todd, M. T., Niv, Y., & Cohen, J. D. (n.d.). Learning to use Working Memory in Partially Observable Environments through Dopaminergic Reinforcement., 8.

Vikbladh, O. M., Meager, M. R., King, J., Blackmon, K., Devinsky, O., Shohamy, D., … Daw, N. D. (2019, May). Hippocampal Contributions to Model-Based Planning and Spatial Memory. Neuron, 102(3), 683–693.e4. Retrieved 2021-07-22, from https://linkinghub.elsevier.com/retrieve/pii/S0896627319301230 doi: 10.1016/j.neuron.2019.02.014

Wagenmakers, E.-J., & Farrell, S. (2004, February). AIC model selection using Akaike weights. Psychonomic Bulletin & Review, 11(1), 192–196. Retrieved 2021-07-07, from http://link.springer.com/10.3758/BF03206482 doi: 10.3758/BF03206482

Wilson, R. C., & Collins, A. G. (2019, November). Ten simple rules for the computational modeling of behavioral data. eLife, 8, e49547. Retrieved 2021-06-28, from https://doi.org/10.7554/eLife.49547 (Publisher: eLife Sciences Publications, Ltd) doi: 10.7554/eLife.49547

Wimmer, G. E., & Shohamy, D. (2012, October). Preference by Association: How Memory Mechanisms in the Hippocampus Bias Decisions. Science, 338(6104), 270–273. Retrieved 2021-07-11, from https://science.sciencemag.org/content/338/6104/270 (Publisher: American Association for the Advancement of Science Section: Report) doi: 10.1126/science.1223252

Xia, L., & Collins, A. G. E. (2020, February). Temporal and state abstractions for efficient learning, transfer and composition in humans. bioRxiv, 2020.02.20.958587. Retrieved 2021-02-08, from https://www.biorxiv.org/content/10.1101/2020.02.20.958587v1 (Publisher: Cold Spring Harbor Laboratory Section: New Results) doi: 10.1101/2020.02.20.958587

